# A mathematical model for velocity-selective arterial spin labeling

**DOI:** 10.1101/2023.08.28.555232

**Authors:** Thomas T. Liu, Eric C. Wong, Divya S. Bolar, Conan Chen, Ryan A. Barnes

**Affiliations:** Center for Functional MRI, University of California San Diego, La Jolla, CA, USA; Department of Radiology, University of California San Diego, La Jolla, CA, USA; Department of Psychiatry, University of California San Diego, La Jolla, CA, USA; Department of Electrical and Computer Engineering, University of California San Diego, La Jolla, CA, USA

**Author notes:** Address correspondence to:* Thomas Liu, UCSD Center for fMRI, 9500 Gilman Drive MC 0677, La Jolla, CA 92093.

**Keywords:** arterial spin labeling, velocity selective, acceleration

## Abstract

**Purpose:** To present a theoretical framework that rigorously defines and analyzes key concepts and quantities for velocity selective arterial spin labeling (VSASL).

**Theory and Methods:** An expression for the VSASL arterial delivery function is derived based on (1) labeling and saturation profiles as a function of velocity and (2) physiologically plausible approximations of changes in acceleration and velocity across the vascular system. The dependence of labeling efficiency on the amplitude and effective bolus width of the arterial delivery function is defined. Factors that affect the effective bolus width are examined, and timing requirements to minimize quantitation errors are derived.

**Results:** The model predicts that a flow-dependent negative bias in the effective bolus width can occur when velocity selective inversion (VSI) is used for the labeling module and velocity selective saturation (VSS) is used for the vascular crushing module. The bias can be minimized by choosing a nominal labeling cutoff velocity that is lower than the nominal cutoff velocity of the vascular crushing module.

**Conclusion:** The elements of the model are specified in a general fashion such that future advances can be readily integrated. The model can facilitate further efforts to understand and characterize the performance of VSASL and provide critical theoretical insights that can be used to design future experiments and develop novel VSASL approaches.

## 1 Introduction

Velocity-selective arterial spin labeling (VSASL) labels blood based on its velocity. While the key features of VSASL have been described in prior work [1, 2, 3], a detailed mathematical model describing the interaction between the labeling process and the velocity profile of the vasculature has been lacking. Furthermore, although there have been qualitative descriptions of the VSASL arterial delivery function [1], a clearly defined quantitative description has not been previously presented. Without a suitable analytical framework, it has been difficult for the field to resolve certain open issues, such as the lack of consensus regarding a “unified generic definition” for cutoff velocity [3]. To address this gap, we present a theoretical framework that rigorously defines and analyzes key concepts and quantities that have been previously been addressed in a largely qualitative fashion. We focus on single module VSASL, which is currently the most widely used implementation of VSASL [3]. A preliminary version of this work was presented in [4].

As the Theory section is rather involved, we start with a guide to the theoretical subsections. After reviewing the basic structure and timing of a typical VSASL scan, we introduce passband and saturation functions in Section 2.1 as representations of the labeling and vascular crushing operations, respectively. We first consider the *matched* case where these functions are implemented with the same pulse sequence module, such as in the initial saturation-based implementation of VSASL [1]. In Section 2.2 we go on to consider the *mismatched* case where the functions are implemented with different pulse sequence modules, such as in recent inversion-based implementations [2], and introduce the notion of an effective saturation function – this Section could be skimmed on an initial reading. We introduce the concept of a boundary velocity in Section 2.3 and build upon it to define the VSASL arterial delivery function in Section 2.4. Adjustments to the definition that take into account arterial blood volume terms are addressed in Section 2.5, but can be passed over on an initial read. In Section 2.6 we incorporate the effects of longitudinal relaxation to derive a general expression for the VSASL magnetization difference as a function of time – this section can serve as a reasonable point to pause and consolidate a basic understanding of the model.

In Sections 2.7 through 2.10, we define and examine quantities that are important for assessing systematic errors in cerebral blood flow (CBF) estimates obtained with VSASL. These include: (i) effective bolus width, (ii) cumulative time integral, (iii) transit delay, and (iv) labeling efficiency. Conditions to achieve the desired effective bolus width are presented in Section 2.11, followed by an examination of bolus width errors due to transit delays and module mismatch in Sections 2.12 and 2.13, respectively. Finally, in Sections 2.14 and 2.15, we introduce adaptations to the model to include spatial variations in transit delays, clarify the contributions of arterial blood volume components, and further refine the conditions required for minimizing bolus width errors. Details of the Bloch simulations and an additional assessment of the mismatch error are then briefly described in the Methods and Results sections, and areas for future work are addressed in the Discussion.

## 2 Theory

We begin by considering a global model in which all functions depend solely on velocity with no explicit dependence on spatial location. Because the key distinguishing feature of VSASL is that the labeling is designed to be largely independent of spatial location, this serves as a natural starting point. We then build upon the insight gained from the global model to consider a local model that incorporates the dependence of velocity on voxel location and size. Expressions are largely derived and presented without relaxation effects, as these can be readily incorporated at a later stage, as shown in Section 2.6. A glossary of key functions and variables is provided in Table 1. Detailed derivations and supplementary material are provided in Supporting Information (SI).

**Table 1:** Glossary of key functions and variables. Functions and variables that have a local model version are indicated by a dependence on the position-dimension variable 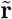 and vessel index *i*.

| Symbol | Description |
| --- | --- |
| $\tau$ | Bolus duration |
| PLD | Post-labeling delay |
| TI | Inflow time |
| VSS | Velocity selection saturation |
| VSI | Velocity selection inversion |
| $v_l, v_{l,\text{eff}}$ | Label/Control module (LCM) nominal and effective cutoff velocities |
| $v_c$ | Vascular crushing module (VCM) cutoff velocity |
| $p(v)$ | Passband function: $p(v) = P_a \cdot p_{\text{ref}}(v)$ |
| $s(v)$ | Saturation function: $s(v) = S_0 \cdot s_{\text{ref}}(v)$ |
| $s_{\text{ref},E}(v)$ | Effective saturation function: $s_{\text{ref},E}(v) = 1 - p(v)/P_a$ |
| $p_{\text{ref}}(v)$ | Reference Passband function with $p_{\text{ref}}(\infty) = 1$ |
| $s_{\text{ref}}(v)$ | Reference Saturation function with $s_{\text{ref}}(0) = 1$ |
| $P_a$ | Steady state value of $p(v)$; with ideal value $P_{a\star} = 1$ for VSS and 2 for VSI |
| $S_0$ | Initial value $s(0)$; with ideal value $S_{0\star} = 1$ |
| $a_a(v)$ | Arterial acceleration function |
| $a_v(v)$ | Venous acceleration function |
| $v_a(t, v_b)$ | Arterial delivery velocity function |
| $v_v(t, v_{\text{cap}})$ | Venous outflow velocity function |
| $v_b, v_{b,i}(\tilde{\mathbf{r}})$ | Boundary velocity |
| $v_{\text{cap}}$ | Mean capillary velocity |
| $V_a$ | Arterial blood volume |
| $c_{a,0}(t, v_b), c_{a,0,i}(\tilde{\mathbf{r}}, t)$ | Arterial delivery function |
| $Q_0, Q_{0,i}(\tilde{\mathbf{r}})$ | Blood flow |
| $\tau_{\text{eff}}$ | Effective bolus width; with ideal value $\tau$ |
| $A_{\text{eff}}$ | Effective bolus area; with ideal value $A_{\text{ideal}} = P_{a\star} \cdot S_{0\star} \cdot \tau$ |
| $\alpha$ | Labeling efficiency; with ideal value $\alpha_{\text{ideal}} = 0.5$ for VSS and 1.0 for VSI |
| $\Delta t$ | Transit delay |

We adopt the naming conventions of [3], where the bolus duration *τ* denotes the time between the labeling/control module (LCM) and the vascular crushing module (VCM), TI (inflow time) denotes the time between the labeling module and the image acquisition module, and PLD denotes the post-labeling delay between the VCM and the image acquisition module. In the labeling condition, the LCM implements either velocity selective saturation (VSS) or inversion (VSI) with module widths of T_VSS_ and T_VSI_, respectively. The VCM typically implements VSS. To simplify the presentation we will assume that both the LCM and VCM act instantaneously such that T_VSS_ = T_VSI_ = 0, and TI = *τ* + T_VSS_ + PLD = *τ* + PLD.

For the global model, we denote *Q*_0_ as the overall blood flow in units of ml/s entering and exiting the cerebrovascular system. The cross-sectional mean blood flow velocity 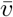 varies along the vascular system, decelerating on the arterial side and accelerating on the venous side. To simplify the notation, we will use the symbol *v* without a bar to denote mean velocity, reserving the 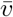 notation to instances where its use provides additional clarity to a derivation. Due to conservation of mass, the overall blood flow at any point in the vascular tree is independent of the mean velocity, i.e. *Q*(*v*) = *Q*_0_.

### 2.1 Passband and saturation functions

We model the creation of VSASL magnetization difference at time *t* = 0 with a unit-less passband function

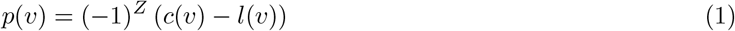

where *c*(*v*) and *l*(*v*) model the normalized actions as a function of velocity *v* of the control and labeling modules, respectively, and *Z* = 0 or 1 for functions *l*(*v*) that aim to perturb spins (saturation or inversion) at velocities above or below, respectively, a specified cutoff velocity [3]. The labeling function *l*(*v*) is either a labeling saturation function *l*_*s*_(*v*) or an inversion function *l*_*i*_(*v*) for VSS and VSI, respectively. The VCM is applied at time *t* = *τ*, and its effects on magnetization are modeled with a saturation function *s*(*v*).

As discussed below, VSASL labeling efficiency depends on both the amplitudes and shapes of the passband and saturation functions. It is thus useful to express the passband function as the product

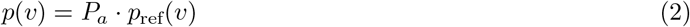

of an amplitude term *P*_*a*_ that represents the steady state value of the passband function and a reference passband function *p*_ref_(*v*) that starts at *p*(0) = 0 and approaches a value of 1.0 for velocities above a labeling cutoff velocity *v*_*l*_. The amplitude term is bounded above by its ideal value *P*_*a*_ *≤ P*_*a⋆*_, where *P*_*a⋆*_ denotes the amplitude of an ideal passband function and is equal to 1.0 and 2.0 for VSS and VSI, respectively. Similarly, the saturation function can be written as the product

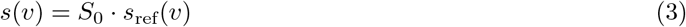

of an amplitude term *S*_0_ = *s*(0) and a reference saturation function *s*_ref_(*v*) that starts at *s*_ref_(0) = 1.0 and approaches zero for velocities above a VCM cutoff velocity *v*_*c*_. The amplitude term is bounded above by its ideal value *S*_0_ *≤ S*_0*⋆*_, where *S*_0*⋆*_ = 1.0 for ideal VCM. A commonly implemented reference saturation function is *s*_ref_(*v*) = sinc(*v/v*_*c*_) [1].

For most VSS implementations, the labeling and VCM saturation functions are implemented with identical pulse sequence modules, so that *l*_*s*_(*v*) = *s*(*v*) = *S*_0_ *·s*_ref_(*v*)). In addition, factors such as *T*_2_ relaxation that reduce the amplitude of the saturation function also reduce the amplitude of the control response, so that *c*(*v*) = *S*_0_ [5]. The resulting passband function is then *p*(*v*) = *S*_0_ *·* (1 *− s*_ref_(*v*)) = *P*_*a*_ *· p*_ref_(*v*) where *P*_*a*_ = *S*_0_ and *p*_ref_(*v*) = (1 *− s*_ref_(*v*)). An example of this for a BIR8 pulse is shown in SI Figure S1.

Example VSS reference passband functions *p*_ref_(*v*) = 1 *− s*_ref_(*v*) and saturation functions *s*_ref_(*v*) (all with *v*_*c*_ = 2.0 cm/s) are shown in Figure 1(a,b). The *s*_ref_(*v*) = sinc(*v/v*_*c*_) function has a soft cutoff and represents a realistic saturation profile under the assumption of laminar flow [1]. In contrast, both the rect function *s*_ref_(*v*) = rect(*v/*(2*v*_*c*_)) and windowed cosine function *s*_ref_(*v*) = cos(*πv/*(2*v*_*c*_)) *·* rect(*v/*(2*v*_*c*_)) have hard cutoffs and cannot be achieved in practice, but are useful for assessing the behavior of ideal profiles with abrupt and smooth transitions, with the windowed cosine serving as a good approximation for the main lobe of the sinc function.

**Figure 1:**
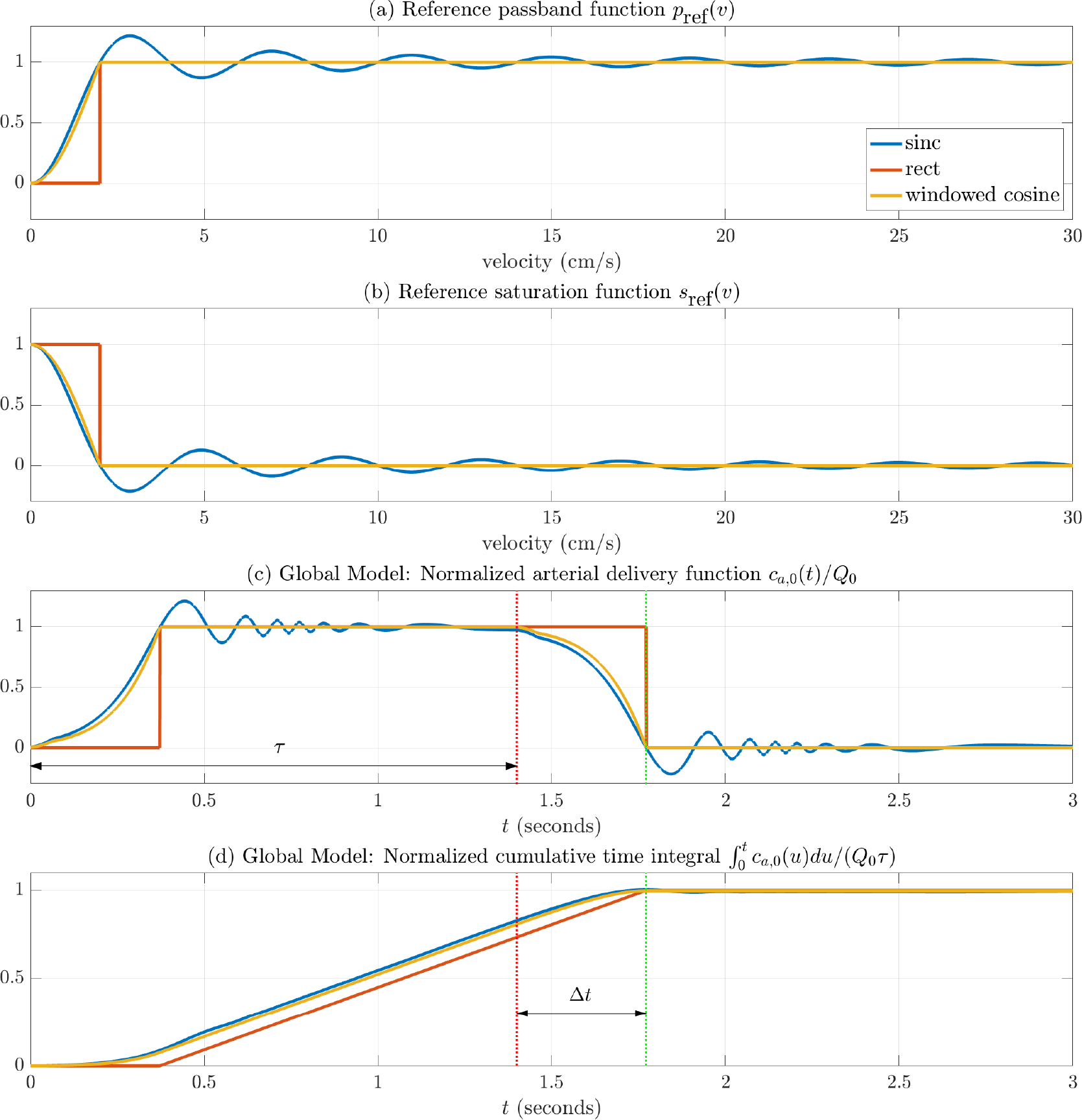
(a,b) Example VSS reference passband *p*_ref_(*v*) = 1 *− s*_ref_(*v*) and saturation *s*_ref_(*v*) functions. Reference saturation functions: sinc(*v/v*_*c*_), rect(*v/*(2*v*_*c*_)), and windowed cosine *s*_ref_(*v*) = cos(*πv/*(2*v*_*c*_)) *·* rect(*v/*(2*v*_*c*_)), all with *v*_*c*_ = 2.0 cm/s. (c) Normalized arterial delivery functions *c*_*a*,0_(*t*)*/Q*_0_ corresponding to the reference passband and saturation functions shown above and assuming *P*_*a*_ = *S*_0_ = 1, *v*_*b*_ = 0.1 cm/s, and *τ* = 1.4 s. (d) Cumulative time integrals 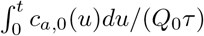, where the ideal value for VSS is 1.0. The red and green dotted vertical lines indicates *t* = *τ* and TI = *τ* + PLD, respectively, with PLD equal to Δ*t*, the transit time required to decelerate from *v*_*c*_ to *v*_*b*_.

### 2.2 Effective reference saturation function

To analyze cases where the LCM and VCM use different modules (e.g. VSI where *l*_*i*_(*v*) *≠ s*(*v*)), it is useful to define an effective reference saturation function as

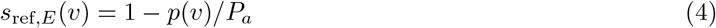

where *s*_ref,*E*_(0) = 1.0 and approaches zero for velocities above the labeling cutoff velocity *v*_*l*_. Note that with this definition we can express the passband function as

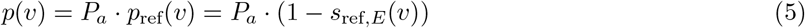

As will be discussed in detail in Section 2.13, mismatches between *s*_ref,*E*_(*v*) and *s*_ref_(*v*) can lead to VSASL quantitation errors. Note when the LCM and VCM use identical modules (as in the VSS examples above), the passband functions are of the form *p*(*v*) = *P*_*a*_ *·* (1 *− s*_ref_(*v*)), so that *s*_ref,*E*_(*v*) = *s*_ref_(*v*) and there is no mismatch (but see also Discussion for cases where implementing a mismatch for VSS may be of interest).

We next consider *s*_ref,*E*_(*v*) for ideal VSI, where the desired passband function is *p*(*v*) = 2*·* (1*−*rect(*v/*(2*v*_*l*_))). This can can be achieved with an ideal control *c*(*v*) = *−*1 and a perfect labeling inversion function *l*_*i*_(*v*) = 1*−*2*·*rect(*v/*(2*v*_*l*_)) that inverts all spins (*l*_*i*_(*v*) = *−*1) for |*v*| *≤ v*_*l*_ while leaving spins with |*v*| *> v*_*l*_ unperturbed (*l*_*i*_(*v*) = 1). Since *p*(*v*) has the form of Eq. 5, we can see from inspection that *s*_ref,*E*_(*v*) = rect(*v/*(2*v*_*l*_)). Note that the perfect labeling inversion function can be written as *l*_*i*_(*v*) = 1 *−* 2 *· s*_ref,*E*_(*v*).

Under the assumption of laminar flow, currently implemented VSI labeling inversion functions can achieve perfect inversion at zero velocity (*l*_*i*_(0) = *−*1.0) but approach an asymptotic value *l*_*i*_(*v*) *→ K* that is less than the desired ideal value of 1.0 for *v > v*_*l*_ [3, 5]. *This behavior can be modeled as l*_*i*_(*v*) = *K −* (1 + *K*)*s*_ref,*E*_(*v*) where |*K*| *≤* 1.0 and *K* = 1 yields the expression for perfect inversion written above. Assuming *c*(*v*) = *−*1, the resulting passband function is *p*(*v*) = (*K* + 1)(1 *− s*_ref,*E*_(*v*)) with amplitude *P*_*a*_ = (*K* + 1) *≤* 2.0. In practice, *s*_ref,*E*_(*v*) is derived with Eq. 4 where *p*(*v*) and *P*_*a*_ are estimated with Bloch simulations or analytical approximations. As with VSS, factors that reduce the value of both the VSI control and inversion functions can be modeled with further reductions in *P*_*a*_.

Figure 2(a,b) shows VSI passband functions *p*(*v*) and effective reference saturation functions *s*_ref,*E*_(*v*) with nominal labeling cutoff velocities of *v*_*l*_ = 1.2 and 2 cm/s using the definition from [3] (see Methods for details on Bloch simulation of the sinc FTVSI module). In this case we can approximate the passband amplitude as *P*_*a*_ = 1.7647 for both responses (see SI Appendix A.8) and compute the effective saturation responses (blue and red curves in panel (b)) using Eq. 4. The VCM saturation response (sinc saturation implemented with BIR8 pulse) with *v*_*c*_ = 2 cm/s (yellow curve) is shown for comparison. Note that there is a considerable mismatch (*s*_ref_(*v*) *≠s*_ref,*E*_(*v*)) between the VCM saturation response and the effective saturation response when the cutoff velocities (*v*_*c*_ = *v*_*l*_ = 2 cm/s) are the same.

**Figure 2:**
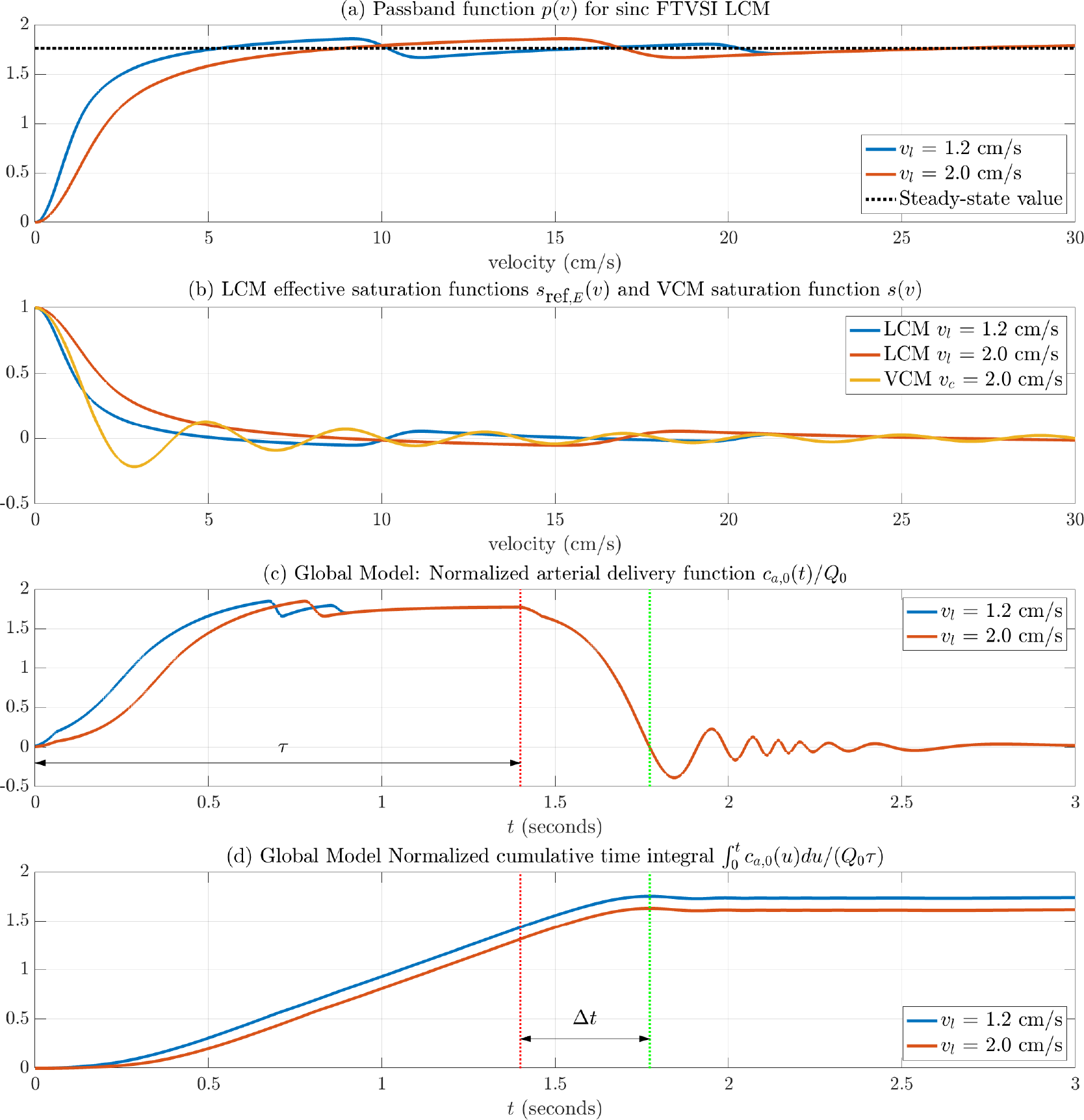
(a) Bloch simulated passband functions *p*(*v*) for sinc FTVSI LCM *v*_*l*_ = 1.2 cm/s (blue) and 2 cm/s (red). The dotted line indicates the steady state value *P*_*a*_. (b) Effective saturation functions computed as *s*_ref,*E*_(*v*) = 1 *− p*(*v*)*/P*_*a*_ functions. Bloch simulated BIR8 VCM saturation function with *v*_*c*_ = 2 cm/s shown for comparison in yellow. (c) Normalized arterial delivery functions *c*_*a*,0_(*t*)*/Q*_0_ corresponding to the passband and saturation functions shown above and assuming *v*_*b*_ = 0.1 cm/s and *τ* = 1.4 s. (d) Cumulative time integrals 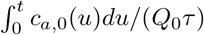, where the ideal value for VSI is 2.0. The red and green dotted vertical lines indicates *t* = *τ* and TI = *τ* + PLD, respectively, with PLD equal to Δ*t*, the transit time required to decelerate from *v*_*c*_ to *v*_*b*_.

### 2.3 Boundary Velocity

In characterizing cerebral perfusion, the goal is to accurately measure the rate of delivery of arterial blood to the capillary beds in brain tissue. To first order, we can say that arterial blood has been delivered to the capillary beds when it has decelerated down to the mean velocity of capillary blood. For the model, we define a boundary velocity *v*_*b*_ such that arterial blood is said to be delivered at time *t* when it has decelerated from its initial velocity *v*(0) to a lower velocity *v*(*t*) = *v*_*b*_. With this definition, the appropriate boundary velocity for the global model is *v*_*b*_ = *v*_*cap*_, where we assume that the mean capillary velocity for normal CBF levels is *v*_*cap*_ = 0.1 cm/s, consistent with the range reported in a recent study [6]. However, higher values of *v*_*b*_ may also be physiologically valid, especially in light of microscopic imaging evidence indicating that up to 50% of oxygen extraction may occur in the parenchymal arterioles [7]. For example, setting *v*_*b*_ = 0.4 would correspond to an arteriole that has a diameter of *∼* 10 microns [8] and represents a point in the arteriolar tree at which substantial oxygen extraction may have already occurred [7]. To simplify the presentation, we will assume *v*_*b*_ = *v*_*cap*_ for the global model, since choosing a lower bound on *v*_*b*_ ensures delivery to the brain tissue. For the local model discussed in Section 2.14, multiple values of *v*_*b*_ can be defined for each voxel, where each value represents the mean velocity of a feeding arteriole at the physical border of the voxel.

### 2.4 Arterial Delivery Function

To derive an expression for the delivery of labeled arterial blood magnetization in the global model, we first consider the magnetization that flows across the boundary velocity *v* = *v*_*b*_ at a time *t ≥* 0, where the LCM is applied at *t* = 0. This corresponds to labeled magnetization *Q*_0_ *· p*(*v*_*a*_(*t,v*_*b*_)) where *v*_*a*_(*t,v*_*b*_) *≥ v*_*b*_ denotes the velocity at labeling time *t* = 0 of arterial blood that will have decelerated to *v*_*b*_ at time *t ≥* 0. As described in SI Appendix A.1, an expression for *v*_*a*_(*t,v*_*b*_) can be derived from the arterial acceleration *a*_*a*_(*v*) as a function of velocity, and an example is shown by the red curve in Figure 3b assuming *v*_*b*_ = *v*_*cap*_. The form of the acceleration function is addressed in SI Section S.1.1.

**Figure 3:**
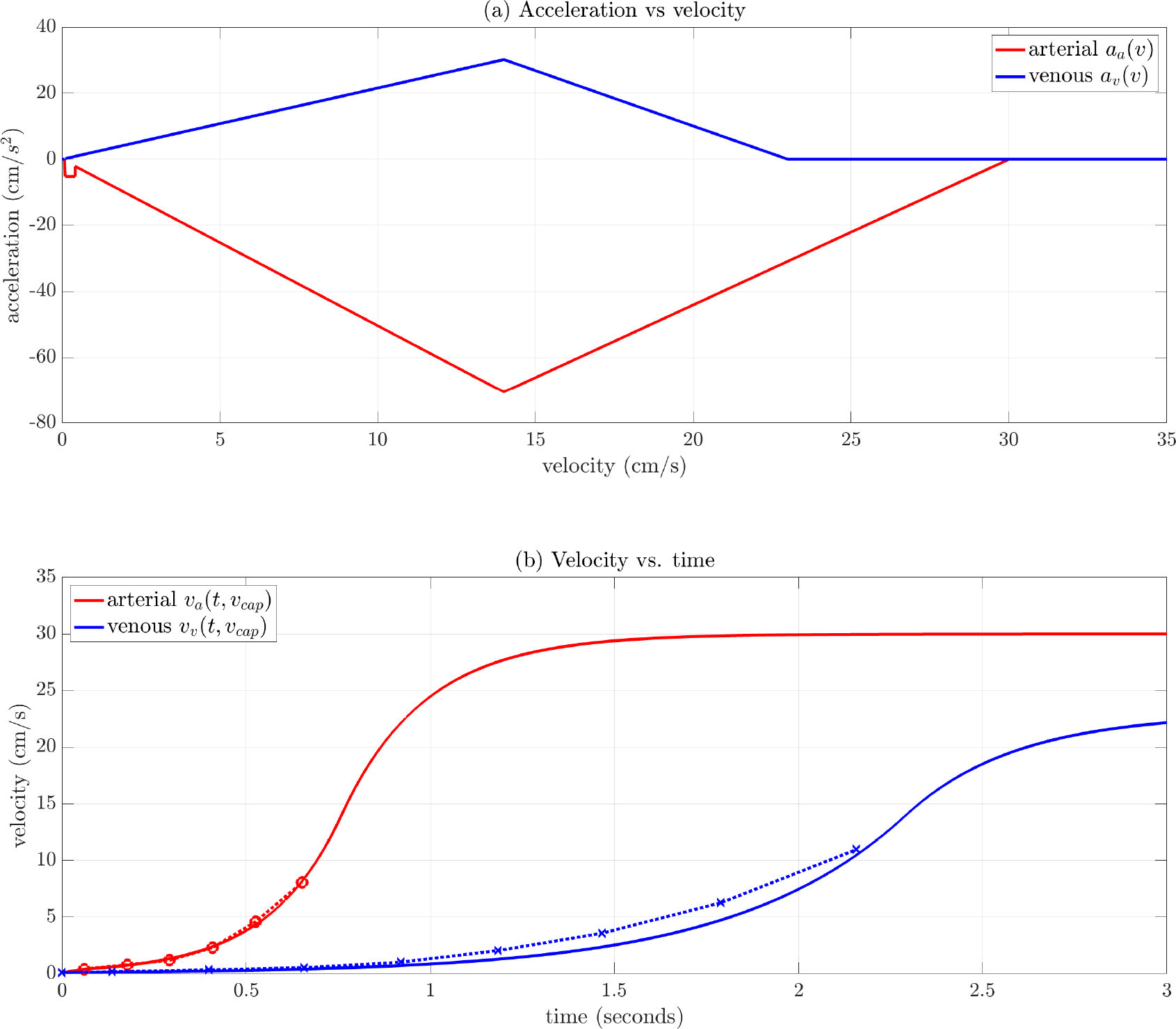
(a) Arterial *a*_*a*_(*v*) (red) and venous *a*_*v*_(*v*) (blue) acceleration functions with parameters described in SI Section S.1.1. (b) Arterial *v*_*a*_(*t, v*_*cap*_) (red) and venous *v*_*v*_(*t, v*_*cap*_) (blue) velocity as a function of time. For the arterial velocity curve, the time axis indicates the arrival time *t* at the boundary velocity of blood with initial velocity *v*_*a*_(*t, v*_*cap*_). For example, blood with an initial velocity *v*_*a*_(1.0, *v*_*cap*_) *≈* 24 cm/s is predicted to decelerate to *v*_*b*_ = *v*_*cap*_ at *t* = 1.0 s. For both panels, the Murray’s law regime applies for velocities below 14 cm/s. In panel (b), symbols and dashed lines show values in this regime from the vascular model adapted from [29].

The VCM saturation applied at *t* = *τ* affects all spins regardless of whether or not they have crossed the velocity boundary. The effect can be modeled by multiplying *Q*_0_ *· p*(*v*_*a*_(*t,v*_*b*_)) by *s*(*v*_*a*_(*τ − t,v*_*b*_)) to yield the arterial delivery function

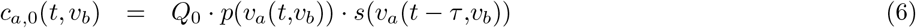

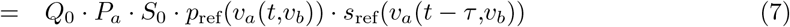

where the 0 in the subscript of *c*_*a*,0_(*t, v*_*b*_) indicates that longitudinal relaxation effects are not included. Note that over the interval [0, *τ*), the term *t − τ <* 0, so that *v*_*a*_(*t − τ,v*_*b*_) *≤ v*_*b*_ represents the velocity at time *τ* of blood that crossed the *v* = *v*_*b*_ threshold at an earlier time *t* and will have therefore decelerated to a velocity less than *v*_*b*_. Thus, for this interval, the multiplication by *s*(*v*_*a*_(*t − τ,v*_*b*_)) represents the saturation at time *τ* of blood that has already crossed the *v* = *v*_*b*_ threshold. Over the interval [*τ*, TI], the term *t − τ ≥* 0, so that *v*_*a*_(*t − τ,v*_*b*_) *≥ v*_*b*_ represents the velocity at time *τ* of blood that will cross the *v* = *v*_*b*_ threshold at a later time *t*. For this interval, multiplication by *s*(*v*_*a*_(*t − τ,v*_*b*_)) represents the saturation at *t* = *τ* of blood that has yet to decelerate to *v*_*b*_. Examples of *v*_*a*_(*t, v*_*b*_) and *v*_*a*_(*t − τ, v*_*b*_) for *v*_*b*_ = *v*_*cap*_ and *v*_*b*_ = 2.0 cm/s are shown in SI Figure S2.

Example arterial delivery functions (normalized by *Q*_0_) are shown in Figure 1c using the reference VSS passband and saturation functions (with *v*_*c*_ = 2 cm/s) from Figure 1(a,b), with *P*_*a*_ = *S*_0_ = 1, *v*_*b*_ = 0.1 cm/s, and *v*_*a*_(*t,v*_*b*_) computed using the acceleration model parameters presented in SI Section S.1.1. Additional examples using the VSI passband functions and sinc saturation function from Figure 2(a,b) are shown in Figure 2c.

### 2.5 Arterial Blood Volume Component

The arterial delivery function accounts for the delivery of arterial blood at velocities *v ≥ v*_*b*_ that has been labeled at *t* = 0. However, when *v*_*b*_ *> v*_*cap*_, it does not account for labeling of arterial blood that can occur when the passband function does not have a hard cutoff and |*p*(*v*)| *>* 0 in the velocity range [*v*_*cap*_, *v*_*b*_]. In addition, it does not account for labeling of blood in the capillary and venous components. These additional terms can be modeled as blood volume components that are initially created by the LCM at *t* = 0 and then modified by the VCM at *t* = *τ* . As shown in SI Appendix A.6, the capillary *V*_*cap*_ and venous *V*_*v*_ volume components are typically negligible.

The volume of arterial blood in the range [*v*_*cap*_, *v*_*b*_] is equal to *Q*_0_ ·Δ*t*_*b*_ where Δ*t*_*b*_ is the time required for blood to decelerate from *v*_*b*_ to *v*_*cap*_ (i.e. *v*_*a*_(Δ*t*_*b*_, *v*_*cap*_) = *v*_*b*_). Note that by definition, the arterial volume term is zero for the global model (since *v*_*b*_ = *v*_*cap*_ and Δ*t*_*b*_ = 0). Thus, we only need to consider it when examining the local model.

Weighting each time increment of blood according to the value of the passband function evaluated at each increment’s initial velocity, we obtain the following expression for the arterial blood volume *V*_*a*_ created by the LCM at *t* = 0:

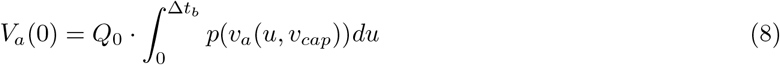

where the variable of integration *u* spans the interval [0, Δ*t*_*b*_], and an example of this integral for *v*_*b*_ = 1 cm/s is indicated by the magenta area in Figure 4a. At *t* = *τ*, the VCM will saturate each time increment of labeled blood volume according to the velocity that it has decelerated to. This can be expressed as

**Figure 4:**
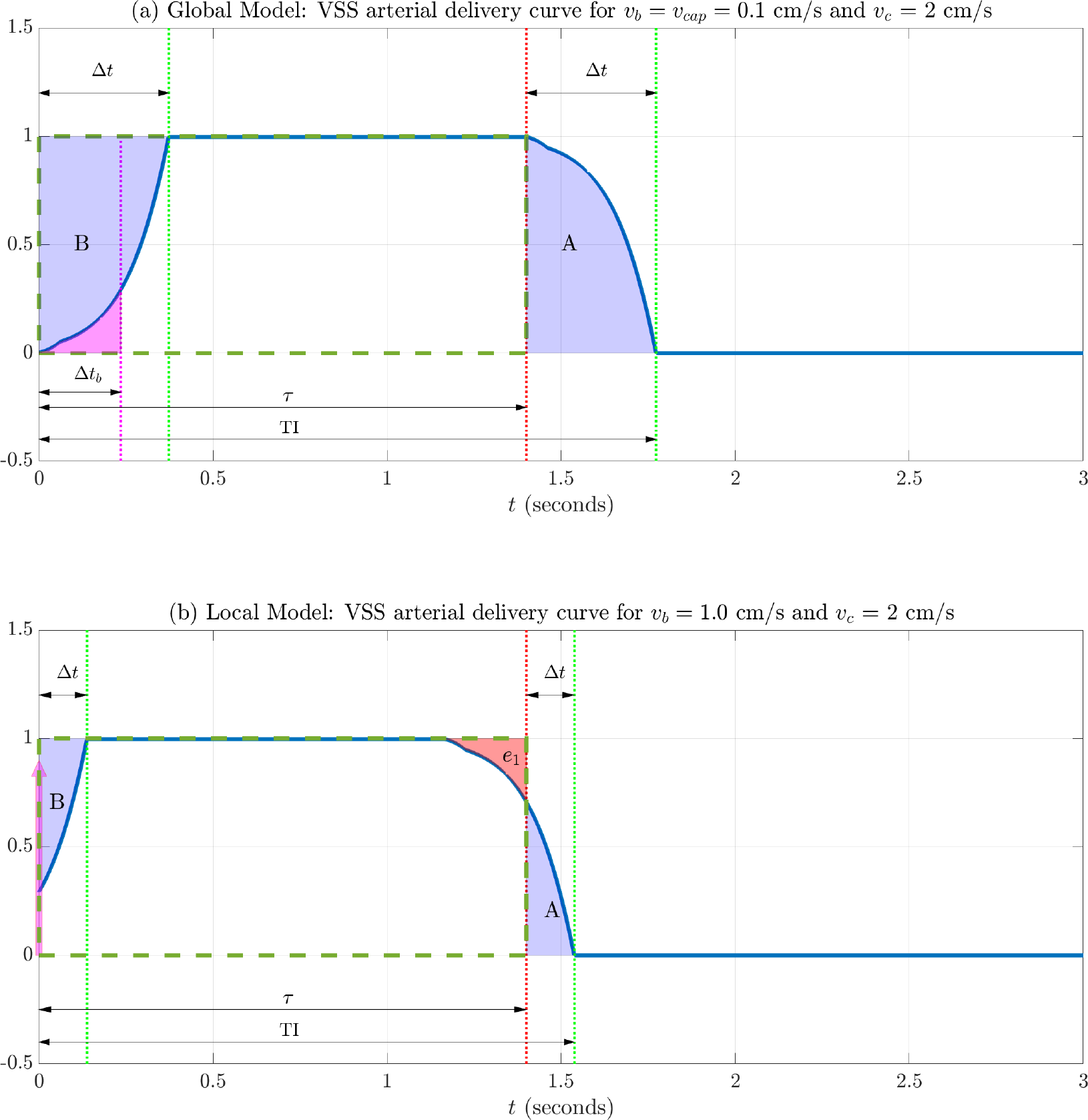
Arterial delivery components for the global and local models, assuming VSS with a windowed cosine saturation function and PLD = Δ*t*. (a) When considering delivery to the capillary beds, the transit delay Δ*t* is the time for blood to decelerate from the *v*_*c*_ = 2 cm/s to *v*_*cap*_ = 0.1 cm/s. As long as PLD *≥* Δ*t*, the leading and trailing edges are complementary and the areas A and B match. As a result, the integral of the arterial delivery function (solid blue curve) is equal to the area of the rectangle (dashed green line) and *τ*_eff_ = *τ* . (b) In the local model, Δ*t* is the time required to decelerate to from *v*_*c*_ = 2 cm/s to *v*_*b*_ = 1 cm/s. As long as PLD *≥* Δ*t*, then areas A and B match. In addition, the error term *e*_1_ is approximately matched by the labeled blood volume component (magenta arrow) injected at *t* = 0, so that the *τ*_eff_ *≈ τ* . This volume component is equal to the magenta area in panel (a), where Δ*t*_*b*_ is the time needed to decelerate from *v*_*b*_ to *v*_*cap*_.

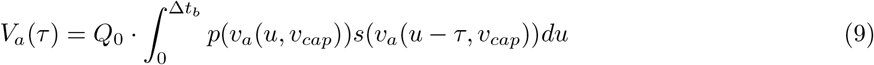

Note that for *τ >* Δ*t*_*b*_, the saturation term *s*(*v*_*a*_(*u − τ, v*_*cap*_)) = *s*(*v*_*cap*_), reflecting the fact that all of the blood in the interval [*v*_*cap*_, *v*_*b*_] has had time to decelerate down to *v*_*cap*_. If this condition is satisfied then

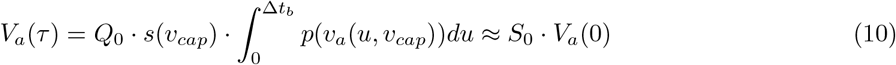

since *s*_ref_(*v*_*cap*_) *≈* 1.0 and therefore *s*(*v*_*cap*_) *≈ S*_0_ for most implementations.

### 2.6 Delivery of Magnetization Difference

Bringing together the terms described in the prior two sections, we can write the overall delivery function without relaxation effects as the sum

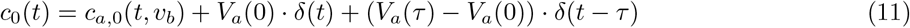

where *δ*(*t*) denotes the Dirac delta function and (i) the term *c*_*a*,0_(*t, v*_*b*_) represents the delivery of arterial blood that starts off at velocities *v ≥ v*_*b*_ and decelerates to *v*_*b*_ at times *t ≥* 0, (ii) the initial arterial blood volume term *V*_*a*_(0) *· δ*(*t*) represents arterial blood in the range [*v*_*cap*_, *v*_*b*_] that is labeled at time *t* = 0 and is not included in the *c*_*a*,0_(*t, v*_*b*_) term, and (iii) the term (*V*_*a*_(*τ*) *− V*_*a*_(0)) *·δ*(*t − τ*) represents the change in the arterial volume term due to the application of the VCM at *t* = *τ* . As noted above, the volume terms are zero for the global model (since *v*_*b*_ = *v*_*cap*_), and thus need to be considered only for the local model (see Section 2.14). In addition, for cases where *s*(*v*) *≈* 1 for *v ∈* [*v*_*cap*_, *v*_*b*_] and *τ >* Δ*t*_*b*_, the VCM has negligible effect on the arterial blood volume term, so that *V*_*a*_(*τ*) *≈ V*_*a*_(0), and thus the volume change term (*V*_*a*_(*τ*) *− V*_*a*_(0)) *· δ*(*t − τ*) is approximately zero.

Incorporating the effects of longitudinal relaxation into the delivery function yields

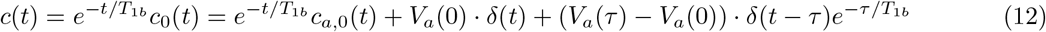

where *T*_1*b*_ denotes the *T*_1_ of blood, and we assume that there is no outflow of labeled blood and a minimal effect of tissue exchange on the relaxation rate, consistent with the assumptions stated in the ASL white paper [9]. Convolution of the delivery function with a residue function 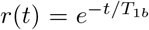 [10] then leads to an expression for the difference in magnetization delivered at time *t*

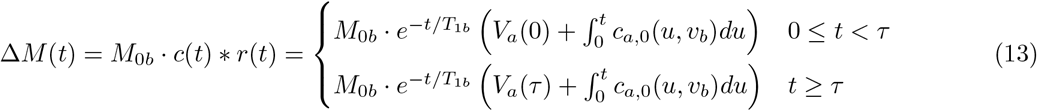

where *M*_0*b*_ denotes the equilibrium magnetization density of arterial blood.

### 2.7 Effective bolus width

For the delivery of an ideal VSASL bolus by some time TI *≥ τ*, it is sufficient to satisfy the following two conditions: (**B1**) 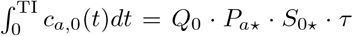 and (**B2**) *V*_a_ (*τ*) = 0, where *P*_*a⋆*_ = 1 or 2 for ideal saturation or inversion, respectively, and *S*_0*⋆*_ = 1.0 for ideal VCM. When these conditions are satisfied, the magnetization difference can be written as 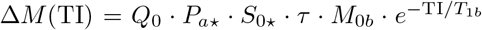 . We first examine Condition B1. As noted above, *V*_*a*_(*τ*) = 0 for the global model, and so we need only consider the impact of volume terms in the local model, which is addressed in Section 2.14.

Note that for Condition B1, the righthand term can be written as *Q*_0_ *· A*_ideal_ where *A*_ideal_ = *P*_*a⋆*_ *· S*_0*⋆*_ *· τ* represents the ideal area under the delivery curve normalized by *Q*_0_. In practice, the actual normalized area under the curve can be written as the effective area

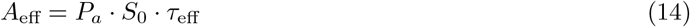

where

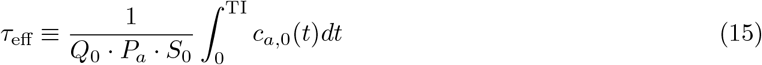

is the **effective bolus width**. With these definitions, we have

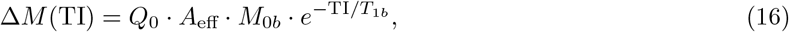

and an estimate of *Q*_0_ can be formed based on measurement of Δ*M* (TI) and estimates of *A*_eff_, *M*_0*b*_ and *T*_1*b*_.

Note that deviations of the effective area *A*_eff_ from the ideal area *A*_ideal_ reflect two effects: (1) the amplitude *P*_*a*_ *·S*_0_ of the normalized delivery function is less than the ideal amplitude *P*_*a⋆*_ *· S*_0*⋆*_, and (2) the effective bolus width *τ*_eff_, which depends on the shape of the normalized delivery function, differs from the desired bolus width *τ* .

### 2.8 Cumulative time integral and transit delay

The effective area *A*_eff_ represents the cumulative normalized signal that been delivered at a specific time *t* = TI. To visualize the dynamic accumulation of signal, it is helpful to also consider the normalized cumulative time integral

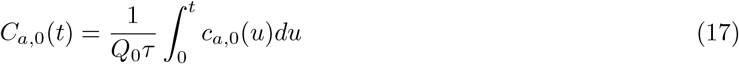

as a function of time *t*. Note that *C*_*a*,0_(TI) = *P*_*a*_ *· S*_0_ *· τ*_eff_*/τ* and thus the ideal value of this quantity is *C*_*a*,0_(TI) = *P*_*a⋆*_ *· S*_0*⋆*_ = 1.0 for VSS and 2.0 for VSI. For the global model where *V*_*a*_(0) = *V*_*a*_(*τ*) = 0, the magnetization difference is related to the normalized cumulative time integral as follows

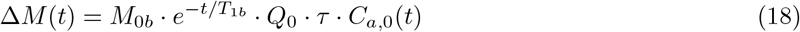

In both Figures 1 and 2, panel (d) shows the normalized cumulative time integrals *C*_*a*,0_(*t*) of the VSS and VSI arterial delivery functions shown in panel (c) of the respective figures, with additional VSI examples shown in SI Figure S4a. In addition, plots of normalized magnetization differences Δ*M* (*t*)*/*(*M*_0*b*_*Q*_0_*τ*) = exp *−t/T*_1*b*_ *· C*_*a*,0_(*t*) for VSS and VSI are shown in SI Figures S3 and S4, respectively. Note that because *v*_*c*_ *> v*_*b*_ in the examples above, the cumulative time integrals reach their steady-state value after some delay following the application of the VCM at time *τ* . This delay is the transit time Δ*t* required for arterial magnetization to decelerate from *v*_*c*_ to *v*_*b*_, and is indicated by the interval between the red (*t* = *τ*) and green (*t* = *τ* + Δ*t*) vertical lines in Figures 1c and 2c.

We can compute the transit delay as follows:

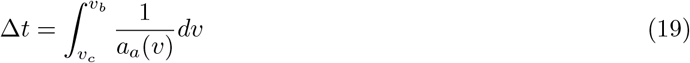

For the above examples, Δ*t* = 372 ms based on the assumed arterial acceleration model parameters for *Q*_0_ = 750 ml/min (see SI Section S.1.1).

At *t* = *τ* + Δ*t* = 1.77, the normalized cumulative time integrals for the rect and cosine tapered profiles both attain the ideal value of 1.0, while the normalized cumulative integral for the sinc profile is very close (1.007) to the ideal value. The approach to and deviation from the ideal value for VSS is further discussed in Section 2.12. In contrast, the VSI cumulative time integrals fall short of reaching the ideal value of 2.0, reflecting two factors: (1) *P*_*a*_ *< P*_*a⋆*_ and (2) a mismatch error that results in *τ*_eff_ *< τ* (see Section 2.13 and Figure 6 for details).

Note that for VSS with matched LCM and VCM (i.e.*l*_*s*_ (*v*) = *s* (*v*)), the transit delay can be viewed as either the time for the leading edge of the arterial delivery function to reach a post-LCM steady state value (e.g. 1.0 for delivery functions in Figure 1(c)) or as the time for the trailing edge to fall to a post-VCM steady state value (typically 0.0). However, this is not the case for the VSI delivery functions in Figure 2(c), due to the use of different modules for the LCM and VCM .Since it is the trailing edge that determines the final approach to steady state of the cumulative time integrals for both VSS and VSI, we have chosen to show the transit delay associated with the trailing edge.

### 2.9 Connection with labeling efficiency

Following the definition from [3], the ideal VSASL labeling efficiency can be written as

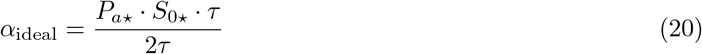

which is equal to 0.5 for perfect VSS and 1.0 for perfect VSI. The actual labeling efficiency is

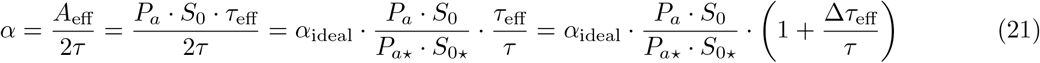

where Δ*τ*_eff_ = *τ*_eff_ *− τ* is the **bolus width error**. Thus, the deviation in labeling efficiency from its ideal value reflects deviations in both the amplitude *P*_*a*_ *· S*_0_ and effective width *τ*_eff_ of the normalized arterial delivery curve from their respective ideal values of *P*_*a⋆*_ *· S*_0*⋆*_ and *τ* .

### 2.10 CBF Estimation Error

As noted below Equation 16, an unbiased CBF estimate can be written as

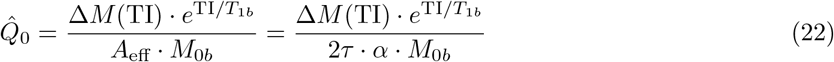

If the estimate uses a value of 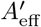 that differs from the actual value *A*_eff_, then the resulting estimate 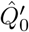 is biased. The fractional error of the biased estimate can be written as

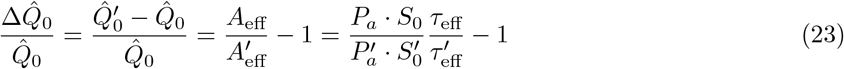

where the primed variables represent the assumed values associated with 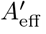. Note that the fractional error may also expressed as 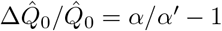. To isolate the error due solely to the bolus width error, we can consider the case where the assumed values of *P*_*a*_ and *S*_0_ are equal to their actual values but the assumed effective bolus width 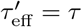 is equal to the ideal bolus width. In this case the resulting fractional error in the CBF estimate

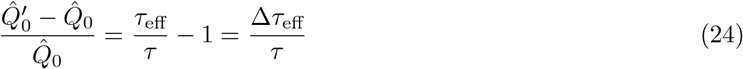

directly corresponds to the fractional bolus width error.

### 2.11 Effective bolus width for matched LCM and VCM

Using Eqs. 5, 7, and 15, the effective bolus width can be written as

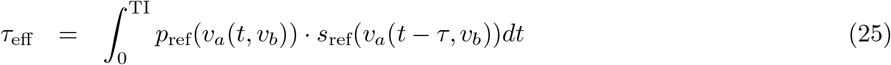

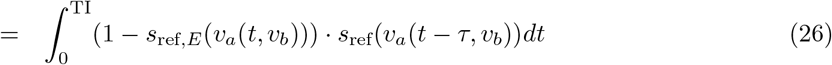

While Eq. 26 can be used to compute *τ*_eff_ for arbitrary functions *s*_ref_(*v*) and *s*_ref,*E*_(*v*), it is also useful to gain some theoretical insight into the conditions needed to achieve *τ*_eff_ = *τ* (and hence satisify condition B1).

We first consider VSS with matched LCM and VCM such that *s*_ref,*E*_(*v*) = *s*_ref_(*v*) and *p*_ref_(*v*) = 1*− s*_ref_(*v*). As shown in SI Appendix A.2, sufficient conditions to obtain *τ*_eff_ = *τ* are as follows:

**Condition W1**: *s*_ref_(*v*) = 0 for |*v*| *≥ v*_*c*_ where the cutoff velocity *v*_*c*_ *≤ v*_*a*_(*τ, v*_*b*_)

**Condition W2**: PLD *≥* Δ*t*,

where the transit delay Δ*t* was defined in Eq. 19.

For typical values of *τ* on the order of one second or more, the term *v*_*a*_(*τ, v*_*b*_) approaches maximum arterial velocities (e.g. 30 cm/s as shown in Figure 3), and Condition W1 can be approached with recommended values of *v*_*c*_ (e.g. 2 cm/s) [3]. Note that implicit in condition W1 is the requirement that Δ*t ≤ τ* . Thus, an alternate viewpoint is that the transit times Δ*t* associated with recommended *v*_*c*_ values are less than typical values of *τ* .

Condition W2 requires that the PLD be chosen greater than the transit time Δ*t*. For example, using the value computed in Section 2.8, the requirement is that PLD *≥* 372 ms for the global model. As discussed below in Section 2.15, transit delays for the local model will typically be smaller than those for the global model, and thus the required PLD will also be smaller.

To gain an intuitive understanding of conditions W1 and W2, it is helpful to consider the normalized VSS arterial delivery curve (using a windowed cosine saturation function) that is depicted by the blue curve in Figure 4a. Note that Condition W1 is strictly satisfied since the windowed cosine is zero for *v > v*_*c*_. As a result, the delivery curve goes to zero at *t* = *τ* + Δ*t*. Furthermore, Condition W2 is satisfied since PLD = Δ*t*, as denoted by the interval between the red (*t* = *τ*) and green (*t* = TI) dotted vertical lines. The effective bolus width is simply the area under the blue curve (since *P*_*a*_ = *S*_0_ = 1.0) over the interval [0, TI]. The desired bolus width *τ* is the area of the dashed green rectangle. When conditions W1 and W2 are met, the leading and trailing edges of the arterial delivery function are complementary, such that the additional area accrued under the trailing edge (area A) compensates for the missing area in the green rectangle (area B), resulting in *τ*_eff_ = *τ* . Note that the importance of the complementary nature of the leading and trailing edges was first noted in [1]. In SI Figure S5a, the arterial delivery curve using a sinc saturation function does not strictly satisfy condition W1 because the sinc function does not have finite support. Nevertheless, due to the oscillating nature of the sinc function, the effective bolus width turns out to be very close to *τ*, as discussed further in the next section.

### 2.12 Errors related to transit delay

When condition W1 is satisfied but W2 is not, then the resulting bolus width error Δ*τ*_eff_ = *τ*_eff_ *− τ* that occurs when PLD *≤* Δ*t* is too small can be considered an error related to the transit delay. As shown in SI Appendix A.3, this error can be written as

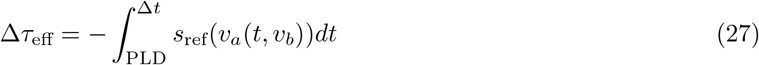

For an ideal *s*(*v*) = rect(*v/*(2*v*_*c*_)), the error can be written as

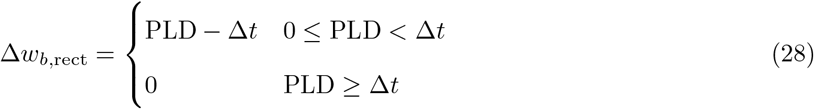

The solid red curve in Figure 5a shows an example of this error term Δ*τ*_eff,rect_ as a function of PLD, assuming *v*_*b*_ = 0.1 cm/s and *v*_*c*_ = 2.0 cm/s, with a normalized version Δ*τ*_eff,rect_*/τ* in panel b. The error curve is negative for PLD *<* Δ*t* and goes to zero when PLD *≥* Δ*t*. Note that for all other saturation functions *s*_ref_(*v*) that satisfy condition W1, the magnitude of the error |Δ*τ*_eff_| *≤* |Δ*τ*_eff,rect_| when PLD *<* Δ*t* is bounded by the error for the rect saturation function (since all valid saturation functions also satisfy |*s*_ref_(*v*)| *≤* 1 for |*v*| *≤ v*_*c*_). As an example, the magnitude of the normalized error for the windowed cosine profile (solid yellow lines in Figure 5) is uniformly smaller than that of the rect profile.

**Figure 5:**
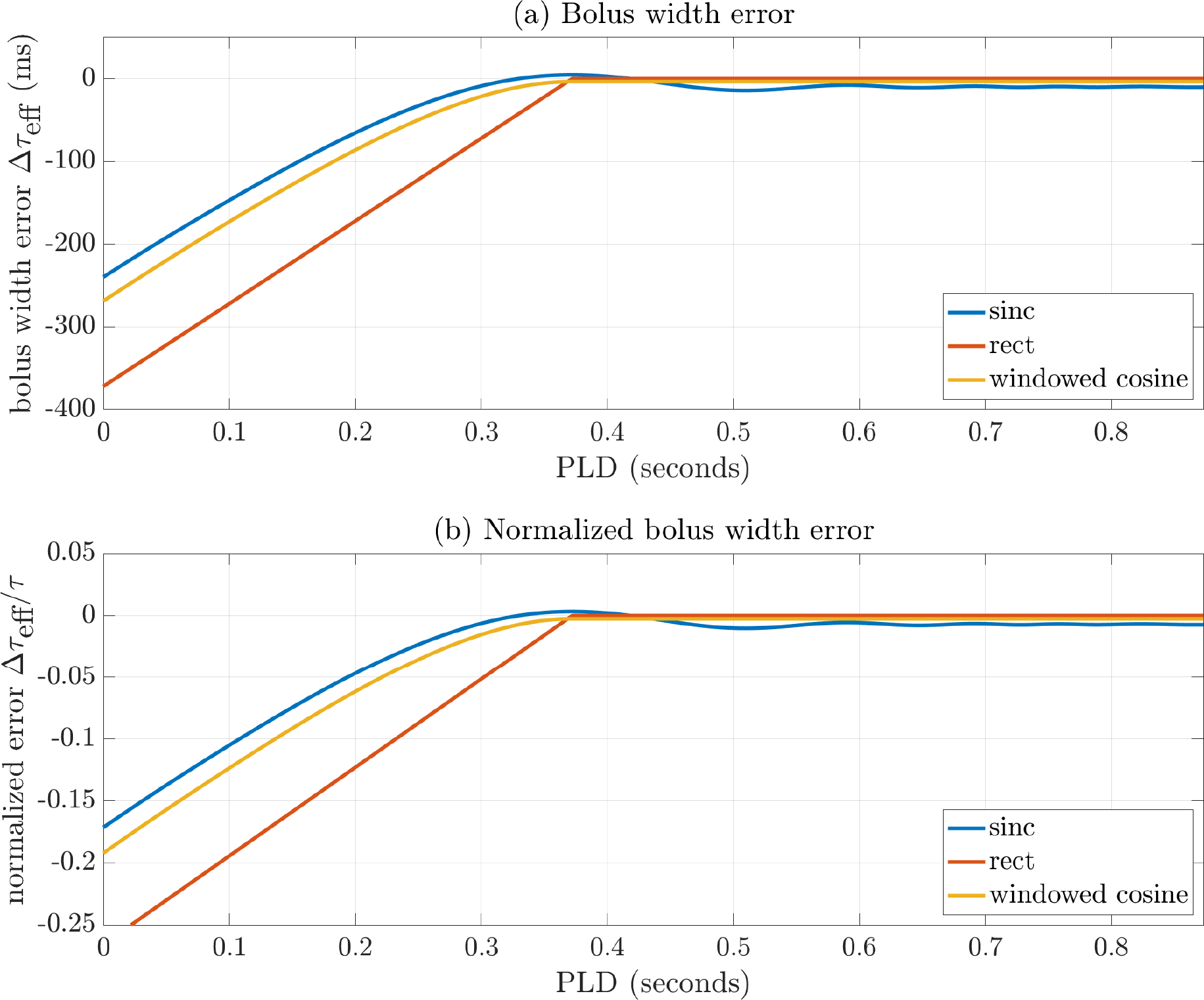
(a) VSS bolus width error Δ*τ*_eff_ and (b) normalized VSS bolus width error Δ*τ*_eff_*/τ* as a function of PLD for sinc (blue), rect (red), and windowed cosine (yellow) profiles assuming *τ* = 1.4 s, *v*_*b*_ = 0.1 cm/s, *v*_*c*_ = 2.0 cm/s, and acceleration function shown in Figure 3.

For *s*_ref_(*v*) = sinc(*v/v*_*c*_), condition W1 can only be approximately satisfied because the sinc function does not have finite support. However, due to the oscillating nature of the sinc function, condition W1 is approximately approached on average, resulting in a normalized bolus width error (solid blue curve in Figure 5b) that is quite small (less than 1%) when PLD *≥* Δ*t*.

### 2.13 Bolus width error due to LCM and VCM mismatch

In Sections 2.11 and 2.12 we considered VSS with matched LCM and VCM, such that *s*_ref,*E*_(*v*) = *s*_ref_(*v*). For current VSI implementations, the LCM and VCM use different pulse sequence modules so that *s*_ref,*E*_(*v*)*≠ s*_ref_(*v*). In addition, as addressed in the Discussion, there may be VSS applications where it may be of interest to have different responses for the LCM and VCM.

As shown in SI Appendix A.4, the bolus width error when there is a mismatch in responses can be written as

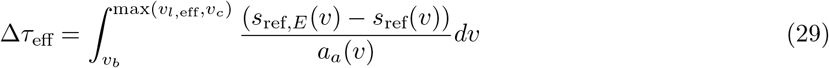

where we introduce the notation *v*_*l*,eff_ to denote the LCM effective cutoff velocity for which *s*_ref,*E*_(*v*_*l*,eff_) *≈* 0. As discussed below, *v*_*l*,eff_ is generally larger than the nominal cutoff velocity *v*_*l*_ for VSI passband functions.

In Figure 2b we showed the effective reference saturation function *s*_ref,*E*_(*v*) corresponding to a VSI passband function *p*(*v*) with *v*_*l*_ = 2 cm/s. Following the convention adopted in prior work [3, 5], the nominal cutoff velocity is defined such that *p*(*v*_*l*_) = 1.0. Applying the definitions from Section 2.1, this corresponds to *p*_ref_(*v*_*l*_) = 1*/P*_*a*_ and *s*_ref,*E*_(*v*_*l*_) = 1 *−* 1*/P*_*a*_, where *P*_*a*_ is the steady-state value of *p*(*v*). For the example VSI passband function, the steady-state value is *P*_*a*_ = 1.7647, resulting in *p*_ref_(*v*_*l*_) = 0.567 and *s*_ref,*E*_(*v*_*l*_) = 0.433. As shown in Figure 2b, *s*_ref,*E*_(*v*) approaches zero at velocities greater than *v*_*l*_, so that *v*_*l*,eff_ *> v*_*l*_. Most current VSI implementations use a VCM with a reference saturation response of the form *s*_ref_(*v*) = sinc(*v/v*_*c*_) with *v*_*c*_ = *v*_*l*_. Reflecting the mismatch (*v*_*l*,eff_ *> v*_*c*_) in the effective cutoff velocities of the LCM and VCM, *s*_ref,*E*_(*v*) has a broader response than *s*_ref_(*v*) (see also Figure 2b for comparison), and the resulting bolus width error Δ*τ*_eff_ in Eq. 29 is negative (since *a*_*a*_(*v*) *<* 0). For the specific parameters and responses shown in Figure 2b, the error is Δ*τ*_eff_ = *−*107 ms (Δ*τ*_eff_*/τ* = *−*7.6%). The negative error reflects the premature saturation by the VCM of labeled spins that have decelerated to the range [*v*_*c*_, *v*_*l*,eff_] at time *t* = *τ* and were otherwise destined to be delivered to the boundary velocity *v*_*b*_ after *t* = *τ* . Essentially, the mismatch in cutoff velocities results in a premature clipping of the bolus such that *τ*_eff_ *< τ* .

The mismatch may be also be viewed as rising from the fact that the leading and trailing edges of the arterial delivery function are not complementary. This lack of complementarity is highlighted in Figure 6a, where the additional area accrued under the trailing edge (area A) is less than missing area in the green rectangle (area B), resulting in *τ*_eff_ *< τ* .

The discussion above suggests that choosing a LCM with *v*_*l*_ *< v*_*c*_ may lead to a better match to the sinc VCM profile. Indeed, Figure 2b shows that the width of *s*_ref,*E*_(*v*) with *v*_*l*_ = 1.2 cm/s is similar to that of *s*_ref_(*v*) with *v*_*c*_ = 2 cm/s. Due to the better matching of the response profiles, the resulting error (Δ*τ*_eff_ = *−*9 ms; Δ*τ*_eff_*/τ* = *−*0.6%)) is negligible. The improved match between the profiles leads to leading and trailing edges that come closer to complementing each other. This is highlighted in Figure 6b, where the additional area accrued under the trailing edge (area A) better matches the missing area in the green rectangle (area B).

**Figure 6:**
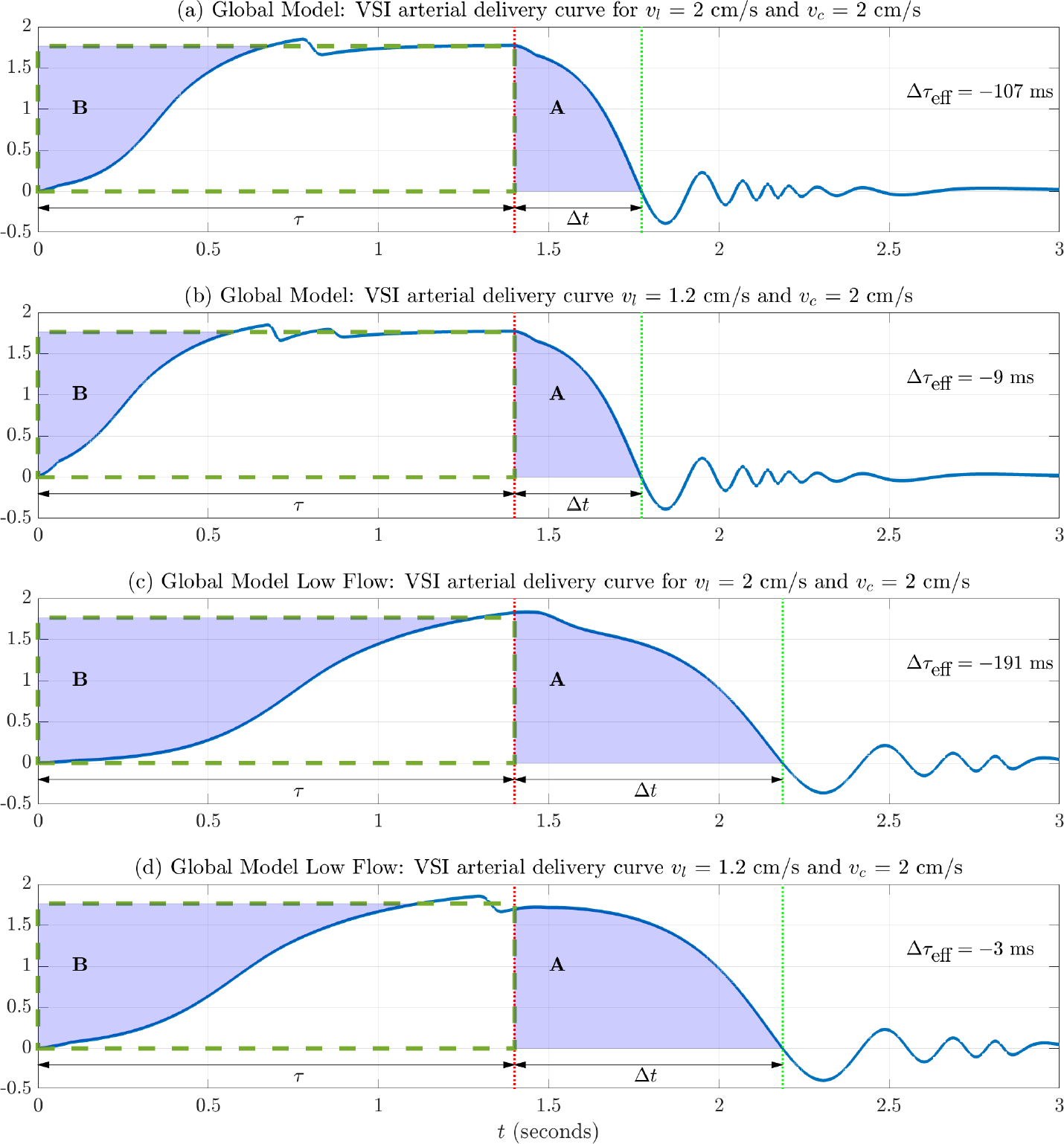
Complementarity of the leading and trailing edges of VSI arterial delivery functions for (a,b) normal flow and (c,d) low flow conditions. (a,c) When *v*_*l*_ = *v*_*c*_ = 2 cm/s, the leading and trailing edges are not complementary. As a result, the additional area (A) accrued under the trailing edge is less than the area (B) excluded by the leading edge, such that the area under the blue curve (up to the green line) is less than the area of the rectangle (dashed green line), resulting in *τ*_eff_ *< τ* and a negative bolus width error Δ*τ*_eff_ *<* 0. The area mismatch and hence the magnitude of the bolus width error is greater for the low flow case. (b,d) When *v*_*l*_ = 1.2 cm/s and *v*_*c*_ = 2 cm/s, the leading and trailing edges nearly complement each other, such that areas A and B are more similar, and the area under the blue curve (up to the green line) is approximately equal to the area of the rectangle (dashed green line), resulting in Δ*τ*_eff_ *≈* 0 for both normal and low flow cases. The red and green dotted vertical lines indicates *t* = *τ* and TI = *τ* + PLD, respectively, with PLD equal to Δ*t*, the transit time required to decelerate from *v*_*c*_ to *v*_*b*_. See text for additional details.

The mismatch examples considered so far have assumed a normal flow condition with global CBF equal to 750 ml/min. In Figure 6(c,d), we consider the complementarity of the leading and trailing edges for a low flow case (460 ml/min) (see Methods for additional low flow parameters). For *v*_*l*_ = *v*_*c*_ = 2 cm/s, the mismatch between the areas A and B is greater than for the normal flow case, leading to a larger error Δ*τ*_eff_ = *−*191 ms (Δ*τ*_eff_*/τ* = *−*13.6%). The larger error reflects the decrease in the magnitude of *a*_*a*_(*v*) that occurs with slower arterial blood flow (see SI Methods and Figure S8a). This decrease leads to an increase in the magnitude of the integrand term 1*/a*_*a*_(*v*) that multiplies the integrand difference term (*s*_ref,*E*_(*v*) *− s*_ref_(*v*)) in Eq. 29. In contrast, for *v*_*l*_ = 1.2 cm/s, there is a relatively good match between areas A and B, resulting in a negligible error Δ*τ*_eff_ = *−*3 ms (Δ*τ*_eff_*/τ* = *−*0.2%), comparable to that observed for the normal flow case. The relative insensitivity to CBF level reflects the fact that changes in the magnitude of *a*_*a*_(*v*) become less important when the saturation profiles are better matched such that the difference term (*s*_ref,*E*_(*v*) *− s*_ref_(*v*)) *≈* 0 is relatively small. Thus, when there is a mismatch in LCM and VCM saturation responses, the bolus width error appears to be more sensitive to CBF level as compared to when the mismatch is minimized. Additional examples of the VSI mismatch error are presented in the Results section.

### 2.14 Local Model

For the local model, we allow for the boundary velocity to vary with the voxel position and also consider multiple feeding arterioles into each voxel, each with its own boundary velocity. Note that the global model may be considered a limiting case of the local model in which all voxels have feeding arterioles with the same boundary velocity *v*_*b*_ = *v*_*cap*_. While this limit cannot be achieved in practice, it could conceivably be approached in the case of extremely high resolution imaging where the dimensions of the voxels becomes small enough that the majority of voxels will only be fed by very small arterioles.

We start by considering a voxel located at position **r** = [*x y z*] with dimensions **L** = [Δ*x* Δ*y* Δ*z*] and defining a position-dimension vector 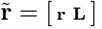, where **L** is independent of position for most cases of interest.

The arterial flow into the voxel can be written as the sum of the flow from 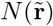 arterial vessels

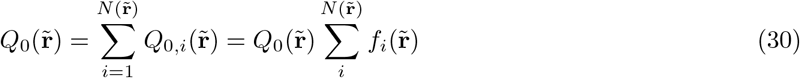

where 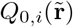 denotes the flow from the *i*th vessel and the fractional flow of the *i*th vessel is defined as 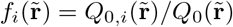. The corresponding labeled inflow delivery function for the *i*th vessel is defined as

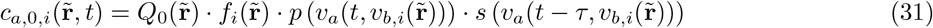

where 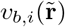 denotes the space and vessel dependent boundary velocity. Summing over the number of vessels yields the overall arterial inflow delivery function for each voxel:

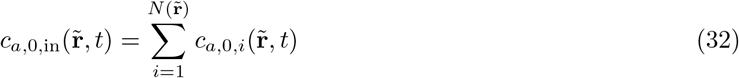

A subtle but important aspect of the local model is that the arterial delivery inflow function does not fully account for the delivery of arterial blood to brain tissue in a voxel. As shown in SI Appendix A.5 for the case of a single feeding arteriole 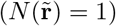, delivery to the capillary bed in a voxel represents the sum of a term that depends on the arterial delivery inflow function and the arterial volume term described in Section 2.5, which accounts for labeled blood that is already in the voxel at *t* = 0. A consequence of this phenomenon is that the PLD requirements for the local model are typically less than those of the global model, since we only need to wait for blood to decelerate to a voxel’s physical boundary velocities, which are generally greater than the capillary velocity. Ignoring macrovascular outflow (which is addressed in the SI), blood that has crossed the voxel’s physical boundaries will ultimately reach the capillary beds. Thus, as long as the PLD is greater than the transit times from *v*_*c*_ to the physical boundary velocities, the VSASL signal will accurately reflect CBF. Examples of this behavior are shown in Figure 4 and SI Figure S5.

### 2.15 PLD Requirements for the Local Model

For an arteriole that ultimately delivers blood to the capillary bed in voxel, the PLD requirement is simply 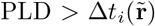 where 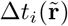 is the time required to decelerate from *v*_*c*_ to the specified 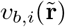. In practice, we will want to set a PLD that satisfies this requirement over a range of anticipated values for 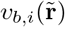. This suggests a requirement of the form:

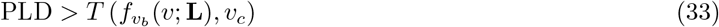

where, for a given value of *v*_*c*_, the function *T* (*·*) (e.g. mean, max, median) returns a summary transit delay based on the probability density function 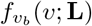 of *v*_*b*_ over all locations 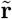 and vessel indices 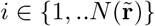 at a given dimension **L**. As addressed further in the Discussion section, we expect that the mean of the density function will decrease as the voxel size decreases. However, further work will be needed to generate suitable approximations of 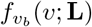, as well to examine the effect of different choices for *T* (*·*).

As an example of the application of Eq. 33, we consider the estimated transit delays shown in Figure 7 for both normal and low flow conditions and three values of *v*_*c*_, where for each value of *v*_*c*_ the transit delays to a range of boundary velocities *v*_*b*_ *∈* [*v*_*cap*_, *v*_*c*_] are plotted. If we assume a uniform distribution of boundary velocities in the interval [1, 2] cm/s and let *T* (*·*) compute the mean transit delay over these boundary velocities, then the resulting requirements for *v*_*c*_ values of 2, 4, and 6 cm/s are that the PLD is greater than 69 ms, 207 ms, and 288 ms, respectively, for the normal flow condition, and greater than 116 ms, 348 ms, and 483 ms, respectively, for the low flow condition.

**Figure 7:**
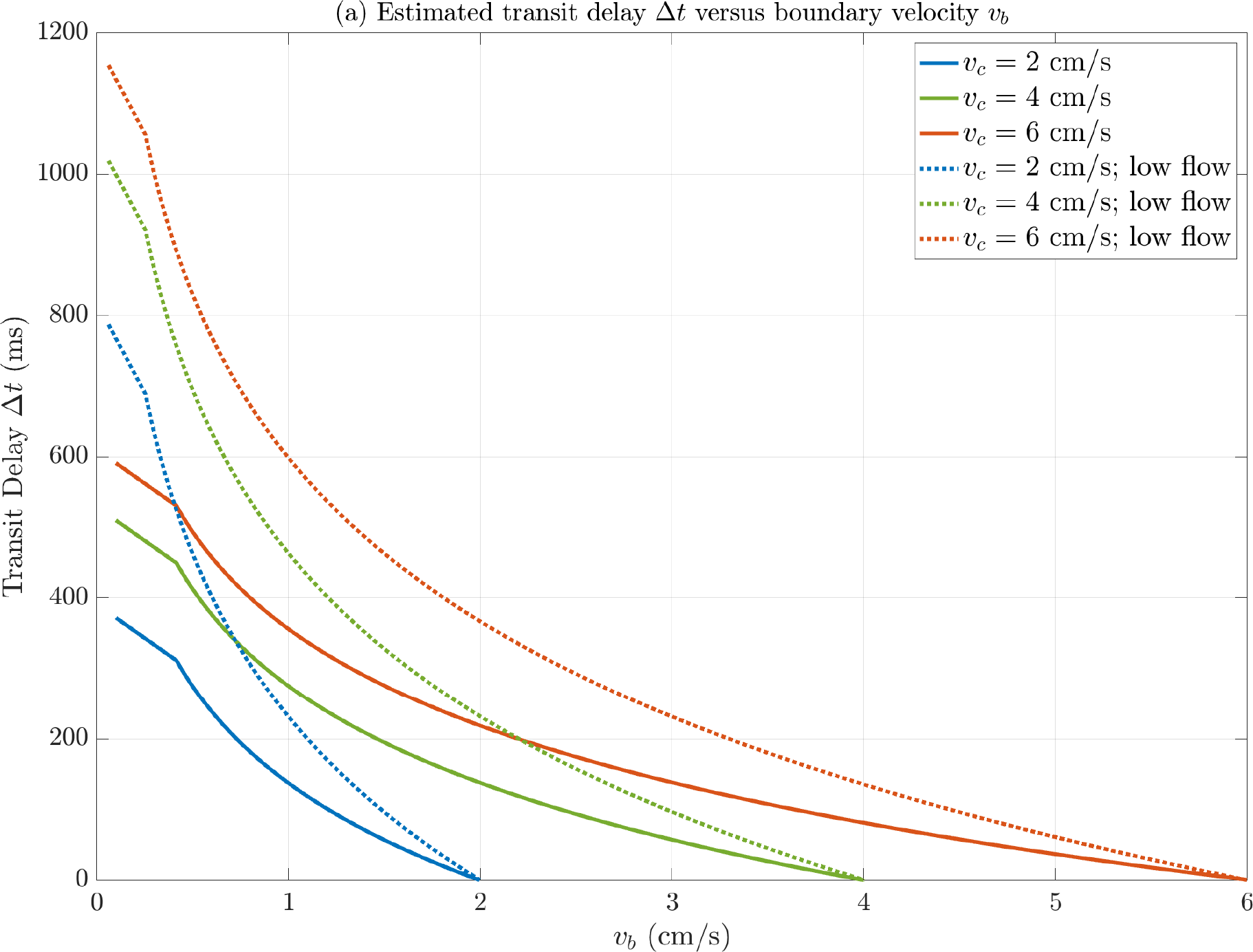
Estimated transit delays versus boundary velocity for normal (solid) and low flow (dot) conditions. For each curve, the transit delays are the times required for arterial blood to decelerate from the specified value of *v*_*c*_ to a range of *v*_*b*_ values, where *v*_*b*_ *∈* [*v*_*cap*_, *v*_*c*_].

## 3 Methods

Estimation of parameters for the acceleration models is described in SI Section S.2. Bloch simulations and laminar flow integration were performed using code from the ISMRM 2022 VSASL Bloch Simulation Tutorial [11]. The BIR8 option with *v*_*c*_ = 2 cm/s was used to simulate the VSS module and the sinc FTVSI option was used to simulate the VSI module with *v*_*l*_ values of 1, 1.2, 2, 2.5, 4, and 6 cm/s. Key parameters were maximum *B*_1_ = 20 *μT*, maximum gradient 50 mT/m, maximum gradient slew rate 150 T/m/s, *T*_1_ and *T*_2_ set to *∞*, and velocity span of 0 to 60 cm/s with a velocity increment of 0.001 cm/s. Arterial delivery functions were obtained using either the BIR8 or sinc FTVSI profiles for the LCM and the BIR8 profile for the VCM. To facilitate evaluating performance over a wider range of *v*_*l*_ values, an empirical approximation for the sinc FTVSI response was derived as described in SI Appendix A.8. This approximation was then used to simulate the LCM VSI responses for *v*_*l*_ ranging from 0.5 cm/s to 6 cm/s with an increment of 0.1 cm/s. For each of the *v*_*l*_ values, arterial delivery functions were obtained using a sinc VCM with *v*_*c*_ values of 2, 4, and 6 cm/s. Delivery functions and associated metrics were computed for both normal and low flow conditions.

## Results

Figure 8 shows bolus width errors for arterial delivery functions computed using the sinc FTVSI approximation LCM for *v*_*l*_ values ranging from 0.5 cm/s to 6 cm/s and sinc VCM with *v*_*c*_ values of 2, 4, and 6 cm/s, with errors for normal and low flow conditions shown in panels (a) and (b), respectively. The errors obtained when *v*_*l*_ = *v*_*c*_ are indicated by the red crosses and range from -57 to -97 ms for normal flow and increased to a range of -169 to -216 ms for the low flow case. Note that the errors obtained when using the FTVSI approximations were slightly different but within 10 ms of those obtained using the Bloch simulations (i.e. plots shown in Figures 2 and 6). For each value of *v*_*c*_, the green x’s indicate the values of *v*_*l*_ that minimize the average error across the normal and low flow cases. These optimal *v*_*l*_ values are 1.2, 2.5, and 4 cm/s for *v*_*c*_ values of 2, 4, and 6 cm/s, respectively, with associated average errors ranging from 2 to 4 ms. Note that for *v*_*c*_ values of 4 and 6 cm/s the low average errors at the optimal values reflect the partial cancelation of small errors with opposite signs for the normal and low flow cases. The VCM sinc saturation profiles are shown by the solid lines in Figure 8c, and for each value of *v*_*c*_ the FTVSI effective saturation profile for the optimal *v*_*l*_ value is shown by a dash-dot line with the same color as the associated sinc profile. In addition, FTVSI effective saturation profiles for *v*_*l*_ = *v*_*c*_ values of 2 and 6 cm/s are shown by the dotted lines (note that *v*_*l*_ = 4 cm/s is also the optimal value for *v*_*c*_ = 6 cm/s and so this curve is not replotted). As discussed above in Section 2.13, the reduction in bolus width error when using an optimal value of *v*_*l*_ reflects the better match between the sinc VCM saturation profile and the FTVSI LCM effective saturation response as compared to the poorer match obtained when using *v*_*l*_ = *v*_*c*_.

**Figure 8:**
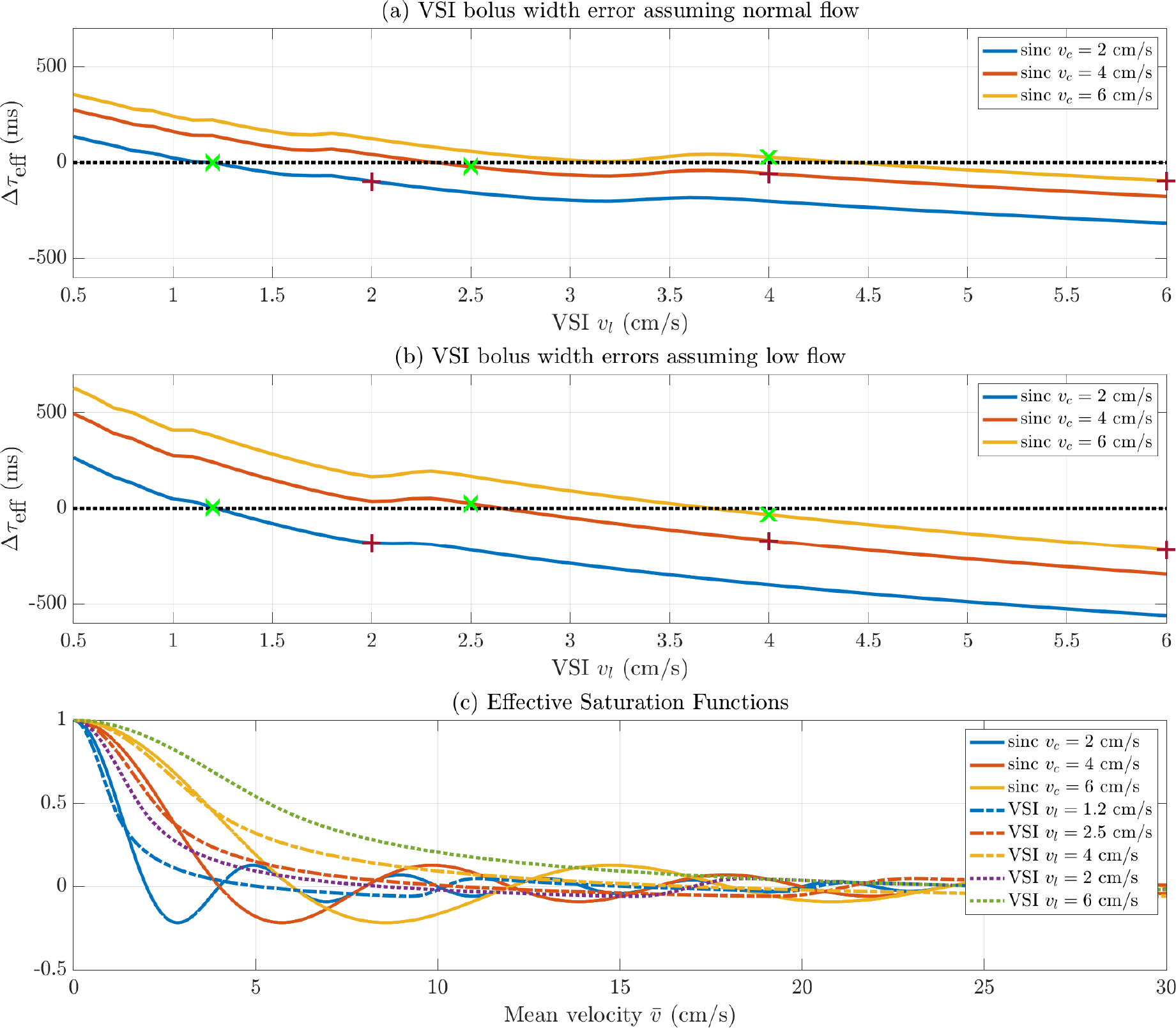
Bolus width errors for (a) normal flow and (b) low flow cases. Each curve shows the bolus width error as a function of FTVSI LCM cutoff velocity *v*_*l*_ for a given value of sinc VCM cutoff velocity *v*_*c*_. Errors obtained when *v*_*l*_ = *v*_*c*_ are indicated by the red crosses. For each *v*_*c*_ curve, the green x’s indicate the error at optimal *v*_*l*_ values that minimize the average error across normal and low flow conditions. (c) Comparison of effective saturation responses. The solid curves show the sinc saturation responses, and the dash-dot curves show the FTVSI effective saturation responses for the corresponding optimal *v*_*l*_ values. The dotted lines show the FTVSI effective saturation responses for values of *v*_*l*_ = *v*_*c*_. Note that *v*_*l*_ = *v*_*c*_ = 4 cm/s is an optimal match for *v*_*c*_ = 6 cm/s, and so is shown only once. Versions of panels a and b that convert the bolus width errors to the percent error in CBF estimates are shown in SI Figure S9.

## 4 Discussion and Conclusion

We have presented a general mathematical model for VSASL that can serve as a framework for both obtaining a deeper understanding of current issues and supporting future analysis efforts. The model rigorously defines the VSASL arterial delivery function, with an explicit description of both blood flow and volume effects. We have used the model to express the dependence of labeling efficiency on both the amplitude and effective bolus width of the arterial delivery function and have examined the factors that affect the effective bolus width. The elements of the model are specified in a general fashion, such that as additional knowledge is gained, these advances may be used to improve the predictive power of the model. For example, in this work we have used simple but physiologically plausible models for arterial and venous acceleration. Future improvements in these acceleration models can be readily integrated into the framework.

Based on the model predictions, we found that there is a potential flow-dependent negative bias in the VSI effective bolus width when setting *v*_*l*_ = *v*_*c*_ with the current definitions of these parameters [3], with the magnitude of the bias becoming larger for lower CBF. We also showed that choice of an optimal *v*_*l*_ *< v*_*c*_ could minimize this bias and make it fairly robust to CBF levels. We introduced the concept of an effective saturation function and demonstrated how a reduction in bias was associated with better matching of the LCM effective saturation function to the VCM saturation function. Future experimental work to validate these theoretical predictions and determine if the negative bias is significant in practice would be of great interest. In addition, since achieving lower *v*_*l*_ values can be challenging for current VSI implementations, the development of alternate approaches for matching the LCM effective saturation and VCM saturation profiles would be of interest.

In most current VSS implementations the LCM and VCM saturation profiles are matched. However, there may be future applications where it is desirable to use saturation profiles with different cutoff velocities, leading to a mismatch between the LCM and VCM profiles and bias in the effective bolus width. It is possible that characterizing the bias over a range of cutoff velocity pairs may provide information about the volume of arterial blood in various velocity ranges.

In the local model version of the current work, we considered delivery of blood to the physical boundary of a voxel and considered a few example cases. Further work to characterize the distribution of boundary velocities as a function of voxel size would be useful for specification of the local model PLD requirement stated in Eq. 33. It likely that the mean boundary velocity (and associated parameters such as mean diameter and number of feeding vessels) will decrease with voxel size. As noted in Section 2.14, in the extremely high resolution limit the boundary velocity of the feeding vessels for the majority of voxels will approach the capillary velocity, in which case the local model converges to the global model. In the present work, we focused on a few example cases (see SI) as pointers to how the framework can be applied to take into account local model features. These cases can serve as the basis for more complicated models that include features such as arterioles that loop around and re-enter a voxel.

One of the advantages of VSASL is its greater insensitivity to vascular transit delays, as compared to spatial ASL approaches. When the cutoff velocity is chosen to be low enough so that the labeling occurs within the voxel, it has been typically assumed that the transit delay is zero [1, 12]. As noted in Section 2.11, one of the requirements for accurate quantification is that PLD *≥* Δ*t*. With the assumption that Δ*t* = 0, we can set PLD = 0, which is the current recommendation of the ISMRM perfusion study group [3]. In this work we explicitly define the transit delay as the time required for arterial blood to decelerate from *v*_*c*_ to *v*_*b*_, and thus we expect there to be non-zero transit delays whenever *v*_*c*_ *> v*_*b*_. Based on the discussion in the previous paragraph, we also expect the transit delay to increase as both the voxel size and mean boundary velocity decreases. While there is some experimental evidence that supports the existence of non-zero transit delays in VSASL [13, 14], more work in this area is warranted. For example, because the predicted transit delays depend on the acceleration profile over the interval [*v*_*b*_, *v*_*c*_], it may be possible to roughly characterize the profile by measuring transit delays over a range of *v*_*c*_ values (where the distribution of *v*_*b*_ values is assumed to not vary across scans within a session when the spatial resolution is fixed). These types of measurements could also potentially be used in future experimental studies to assess how the acceleration profile and other parameters of the model (such as the distribution of boundary velocities) might vary across conditions, including aging and cerebrovascular diseases such as Moyamoya.

In our model we have used simple acceleration models that relate mean acceleration to mean velocity. These simple models assume a monotonic decrease in mean velocity as blood enters through the arterial tree and a monotonic increase as blood exits through the venous tree. As discussed in SI Sections S.1.1 and S.2, the general form of the simple model is well supported by prior experimental and theoretical work. However, it important to keep in mind that detailed studies of flow patterns in realistic vessels [15, 16, 17], while relatively sparse, suggest that velocity profiles are extremely complex, especially at bifurcations, where it likely that deceleration and acceleration may be nonlinear. In addition, velocity profiles have been shown to exhibit a dependence on cardiac phase [17, 18, 19], with significant deviations from laminar flow profiles. Finally, since most current VSASL implementations use a single velocity encoding gradient direction [3], the effective labeling efficiency may vary across the length of a vessel as the angle to the gradient changes, leading to non-monotonic changes in the labeled velocity component along the vascular tree. While these effects may largely average out over space and time for a typical VSASL experiment, a greater understanding of the effects would be of interest for future work. Furthermore, it is likely that characterizing the spatiotemporal complexity of flow patterns may be critical for developing a model for acceleration-sensitive ASL [20].

Following prior work [3, 1], we have assumed a laminar flow profile for the computation of the passband *p*(*v*) and saturation *s*(*v*) functions. We have then assumed that the magnetization difference that is delivered as a function of time is described by the time required to move from a given initial mean velocity (e.g. *v*_*a*_(*t, v*_*b*_)) to the boundary velocity *v*_*b*_. However, this assumption does not take into account the variation in transit times that may be experienced by blood at different radial locations within a vessel. Prior work examining dispersion effects in spatial ASL [21, 22, 23, 24, 25] suggests that the variation in transit times across layers due to laminar flow may be largely confined to the larger vessels followed by “plug-flow-like” transit times in the smaller vessels, possibly reflecting rapid mixing across layers and the complex flow effects described in the previous paragraph. SI Appendix A.7 sketches out a general approach for extending the definition of the passband function to incorporate distributions describing the dependence of transit times on radial position and vessel mean velocity. This extended approach requires detailed modeling of microvascular flow and parameters that may not yet be available in the literature. Further work will be needed to determine if the extended approach offers significant advantages over the simple approach used in this paper, especially given the likelihood that the simple approach offers a reasonable first order approximation of average behavior over the range of parameter distributions describing the microvascular system.

In this paper we have focused on single module VSASL, which is currently the mostly widely used implementation. Recent work has shown that dual module VSASL approaches can offer significant gains in performance [26, 27]. As compared to single module VSASL, the creation of the magnetization difference in dual module VSASL is more complicated and does not lend itself to a description by the passband function used in this work. Future extensions of the model to describe dual module VSASL would be of interest, especially if the dual module approach gains more widespread adoption.

The definition of the arterial delivery function in Eq. 7 implicitly assumes that the application of the VCM determines the end of the arterial bolus. Since the maximum available bolus width may be lower in physiological states with high velocity or cases where there is limited spatial coverage of the VSASL labeling pulse, the available bolus width may be less than *τ* [28]. Extensions of the current model to model this effect would be of interest. This would most likely entail introducing a joint dependence on velocity and space into the passband function that would allow for an estimation of the physical bolus width.

In conclusion, we have introduced a mathematical model that clearly and rigorously defines key concepts in VSASL that have previously been addressed in a largely qualitative fashion. We anticipate that by providing a fundamental framework for analyzing VSASL, this work will greatly facilitate further efforts to understand and characterize the performance of VSASL and provide critical theoretical insights that can be used to design future experiments and develop novel VSASL approaches.

## 5 Data Availability Statement

Analysis code and files to generate the figures and results presented in this paper will be made available upon publication through the Open Science Framework DOI:10.17605/OSF.IO/WKM54.

## SI Figure Captions

**Figure S1.**(a) Bloch simulated BIR8 label and control profiles with either *T*_1_ = *T*_2_ = *∞* for reference profiles (*l*_ref_(*v*) and *c*_ref_(*v*)) and *T*_1_ = 1660 ms and *T*_2_ = 150 ms for *l*(*v*) and *c*(*v*). As compared to the reference profiles, *l*(*v*) and *c*(*v*) are both scaled by a factor of *P*_*a*_ = 0.92 (see Methods for additional simulation parameters). (b) Passband functions *p*_ref_(*v*) = *c*_ref_(*v*) *− l*_ref_(*v*) and *p*(*v*) = *c*(*v*) *− l*(*v*) corresponding to the label and control profiles shown in panel (a). Note that *p*(*v*) = *P*_*a*_ *·p*_ref_(*v*) as shown by the matching of the red solid curve and yellow dotted curve.

**Figure S2.**Plots of *v*_*a*_(*t, v*_*b*_) (solid) and *v*_*a*_(*t − τ, v*_*b*_) (dash) for *v*_*b*_ = *v*_*cap*_ = 0.1 cm/s (red) and *v*_*b*_ = 2.0 cm/s (magenta) calculated using the arterial acceleration model parameters from Section S.1.1. The black vertical line indicates *t* = *τ* . At *t* = *τ*, we have *v*_*a*_(*t − τ, v*_*b*_) = *v*_*a*_(0, *v*_*b*_) = *v*_*b*_, and hence the intersection of the *v*_*a*_(*t − τ, v*_*b*_) dashed curves with the black vertical line occur at the points (*τ, v*_*b*_). For *v*_*b*_ = 2.0 cm/s, this point is indicated by the intersection of the black vertical line with the horizontal magenta line. For times *t ≤ τ*, the argument *t − τ* is negative, resulting in *v*_*a*_(*t − τ, v*_*b*_) taking on values less than or equal to *v*_*b*_.

**Figure S3.**Plots of normalized magnetization difference 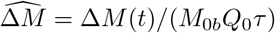 for VSS with sinc (blue), rect (red), and windowed cosine (yellow) saturation functions and *P*_*a*_ = *S*_0_ = 1. The dotted purple line shows the exponential decay curve exp(*−t/T*_1*b*_) with *T*_1*b*_ = 1660 ms. Note that the 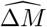 curves can be written as the product 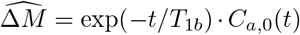 of the exponential decay curve and the normalized cumulative time integral *C*_*a*,0_(*t*) curves shown in Figure 1d. Since the *C*_*a*,0_(*t*) curves approach their steady-state value of 1.0 at *t* = *τ* + Δ*t*, the 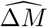 curves follow the exponential decay curve exp(*−t/T*_1*b*_) for *t ≥ τ* + Δ*t*.

**Figure S4.**(a) Normalized cumulative time integrals *C*_*a*,0_(*t*) for VSI with *v*_*l*_ = 1.2 cm/s and *v*_*l*_ = 2 cm/s for both normal (red, blue) and low flow conditions (yellow, purple), with *S*_0_ = 1.0 and *P*_*a*_ = 1.7647 (black dotted line). (b) Plots of normalized magnetization difference 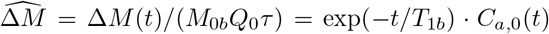 The dotted black line shows the curve *P*_*a*_ *·* exp(*−t/T*_1*b*_) where *T*_1*b*_ = 1660 ms. Note that the *C*_*a*,0_(*t*) curves for *v*_*l*_ = 1.2 cm/s approach the steady-state value of *P*_*a*_ at *t* = *τ* + Δ*t*_*normal*_ and *t* = *τ* + Δ*t*_*low*_ for normal and low flow conditions, respectively. As a result, the 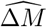 curves for *v*_*l*_ = 1.2 approximately follow the exponential decay curve *P*_*a*_ *·* exp(*−t/T*_1*b*_) for *t ≥ τ* + Δ*t*_*normal*_ and *t ≥ τ* + Δ*t*_*low*_ for normal and low flow conditions, respectively. The lower relative amplitudes of both the *C*_*a*,0_(*t*) and 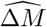 curves for *v*_*l*_ = 2.0 cm/s reflect the effects of the mismatch discussed in Section 2.13.

**Figure S5.**Arterial delivery components for the global and local models, assuming VSS with a sinc saturation function. (a) When considering delivery to the capillary beds, the transit delay Δ*t* is the time for blood to decelerate from the *v*_*c*_ = 2 cm/s to *v*_*cap*_ = 0.1 cm/s. As long as PLD *≥* Δ*t*, the leading and trailing edges are complementary and the areas A and B match. As a result, the integral of the arterial delivery function (solid blue curve) over the interval [0, TI] is equal to the area of the rectangle (dashed green line) and *τ*_eff_ = *τ* . (b) In the local model, Δ*t* is the time required to decelerate to from *v*_*c*_ = 2 cm/s to *v*_*b*_ = 1 cm/s. As long as PLD *≥* Δ*t*, then areas A and B match. In addition, the error term *e*_1_ is approximately matched by the labeled blood volume component (magenta arrow) injected at *t* = 0, so that the *τ*_eff_ *≈ τ* . This volume component is equal to the magenta area in panel (a), where Δ*t*_*b*_ is the time needed to decelerate from *v*_*b*_ to *v*_*cap*_.

**Figure S6.**Arterial delivery functions (panels a, c, e) and cumulative delivery signals (panels b, d, f) for the local model case examples, assuming sinc VSS with *v*_*c*_ = 2 cm/s. Note that the arterial delivery functions are normalized by 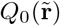, and the cumulative delivery signals include the effect of an initial arterial volume term for vessels that deliver blood to the capillary bed. Case 1 (panels a, b): two arterioles with different boundary velocities (0.4 cm/s and 1.5 cm/s) deliver blood to the capillary bed. Case 2 (panels c, d): an arteriole passes through the voxel with entry and exit velocities of 1.5 cm/s and 0.4 cm/s, respectively. Case 3 (panels e,f): arteriole that branches into two child arterioles, with one delivering blood to the capillary bed and the other exiting the voxel. See text for additional details.

**Figure S7.**Arterial delivery functions (panels a, c, e) and cumulative delivery signals (panels b, d, f) for the local model case examples, assuming windowed cosine VSS with *v*_*c*_ = 2 cm/s. Note that the arterial delivery functions are normalized by 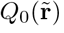, and the cumulative delivery signals include the effect of an initial arterial volume term for vessels that deliver blood to the capillary bed. Case 1 (panels a, b): two arterioles with different boundary velocities (0.4 cm/s and 1.5 cm/s) deliver blood to the capillary bed. Case 2 (panels c, d): an arteriole passes through the voxel with entry and exit velocities of 1.5 cm/s and 0.4 cm/s, respectively. Case 3 (panels e, f): arteriole that branches into two child arterioles, with one delivering blood to the capillary bed and the other exiting the voxel. See text for additional details.

**Figure S8.**(a) Arterial *a*_*a*_(*v*) (red, brown) and venous *a*_*v*_(*v*) (blue, light blue) acceleration functions with parameters described in Section S.1.1 for normal (solid) and low flow (dash) conditions. (b) Arterial *v*_*a*_(*t, v*_*cap*_) (red, brown) and venous *v*_*v*_(*t, v*_*cap*_) (blue, light blue) velocity as a function of time for normal (solid) and low flow (dash) conditions). For both panels, the Murray’s law regime applies for velocities below 14 cm/s. In panel (b), symbols and dashed lines show values in this regime from the vascular model adapted from [29].

**Figure S9.**Percent error in CBF estimates for (a) normal flow and (b) low flow cases, obtained with Equation 24 using bolus width errors from Figure 8 and *τ* = 1.4 s. Each curve shows the percent error as a function of FTVSI LCM cutoff velocity *v*_*l*_ for a given value of sinc VCM cutoff velocity *v*_*c*_. Errors obtained when *v*_*l*_ = *v*_*c*_ are indicated by the red crosses. For each *v*_*c*_ curve, the green x’s indicate the error at optimal *v*_*l*_ values that minimize the average error across normal and low flow conditions.

**Figure S10.**Approximations to the VSI passband functions. (a) Solid lines show Bloch simulated VSI passband functions for *v*_*l*_ = 2.0 cm/s and *v*_*l*_ = 1.2 cm/s prior to laminar flow integration, while the dotted lines show the corresponding approximations from Eq. A.19. (b) Differences between the simulated passband functions and the approximations. (c) Solid lines show Bloch simulated VSI passband functions after laminar flow integration, while dotted lines show corresponding approximations from Eq. A.20. (d) Differences between the simulated passband functions and the approximations with laminar flow integration. Note that the velocity range for the top two rows is double that of the bottom two rows, and the errors shown in (d) are an order of magnitude less than those in (b).

