## Supporting Information for "A mathematical model for velocity-selective arterial spin labeling"

October 3, 2023

*Running Head:* Velocity-selective ASL Model

*Address correspondence to:*

Thomas Liu, UCSD Center for fMRI, 9500 Gilman Drive MC 0677, La Jolla, CA 92093

Supporting Information submitted to Magnetic Resonance in Medicine

#### S.1 Supporting Theory

##### S.1.1 Acceleration Models

In this section we present simple expressions for $a_a(v)$ and $a_v(v)$ that can be used to demonstrate the computation of the delivery function and associated terms described in prior sections and in the Supporting Information (SI) Appendix. The expressions are intended to capture the key aspects of the average arterial deceleration and venous acceleration along the cerebrovascular system.

To first order, the vessels in the cerebrovascular system can be divided into two major regimes: (1) a Murray's law regime for smaller vessels, in which flow $Q \propto r^3$ is proportional to the third power of vessel radius $r$ and (2) a da Vinci's rule regime for larger vessels, in which flow $Q \propto r^2$ is proportional to the second power of vessel radius [1, 2]. In the Murray's law regime, the relation $Q \propto r^3$ requires that the vessel radii at each branching point satisfy the relation $r_k^3 = \sum_{i=1}^{n} r_{k+1,i}^3$ where $k$ and $k+1$ index the branch levels of the parent and child arterial vessels, respectively, $i$ indexes the child vessels, and the branching ratio $n$ denotes the number of child vessels at each levels [3, 4]. With the simplifying assumptions of (a) constant branching ratio $n$ across all levels and (b) equal radii across child vessels at each level, we can write $r_{k+1}/r_k = n^{-1/3}$. Furthermore, because $Q \propto A_c v$, where $A_c$ denotes vessel cross-sectional area, the vessel velocity $v \propto r$ is proportional to radius [3, 4] and therefore $v_{k+1}/v_k = n^{-1/3}$ . With the additional assumption that the vascular network exhibits a space-filling fractal branching pattern, the ratio of vessel lengths follows $l_{k+1}/l_k = n^{-1/3}$ [1]. Combining the last two relations leads to the conclusion that the average time spent in each vessel $\Delta t_k = l_k/v_k = \lambda$ is a constant that is independent of the branch level $k$. Denoting $v_0$ as the velocity at the $k = 0$ branch level and noting that the cumulative time spent flowing in vessels up to the start of the $k$th level is $t_k = k\lambda$, we can write $v_k = v_0 n^{-k/3} = v_0 n^{-t_k/(3\lambda)} = v_0 e^{-t_k \log n/(3\lambda)}$, which has the form $v \propto e^{-\kappa_a t}$, where $\kappa_a > 0$ is a scaling parameter with units of $s^{-1}$. As a consequence of the exponential form for velocity, the mean arterial acceleration can be expressed as $a_a(v) = -\kappa_a v$ in the Murray's law regime. Similar arguments apply to the venous side, resulting in $a_v(v) = \kappa_v v$ with $\kappa_v > 0$.

In the da Vinci's rule regime, the relations $Q \propto r^2$ and $Q \propto A_c v$ imply that velocity is independent of vessel radius, and thus in this regime the mean acceleration is zero (i.e. $a_a(v) = a_v(v) = 0$). Based upon experimental and theoretical considerations, the transition between regimes is estimated to start at vessel diameters around 0.4 to 0.6 mm [1, 4, 5], reaching the da Vinci's rule regime at vessel diameters around 0.6 to 0.7 cm (e.g. in the common carotid artery) [6]. In our simple model for $a_a(v)$, we initially include four zones: (1) zero mean acceleration for velocities below the mean capillary velocity $v_{cap}$, (2) Murray's law regime for velocities between $v_{cap}$ and an arterial transition velocity $v_{t,a}$, (3) a linear transition between Murray's law and da Vinci's rule regimes starting at $v_{t,a}$ and ending at the maximum arterial velocity $v_{\max,a}$; and (4) da Vinci's rule regime for velocities above $v_{\max,a}$. In addition, to provide a better empirical fit to prior experimental data (see SI Methods), we allow for a fifth zone in which there is a constant negative acceleration $-a_0$ for a small range of velocities $v \in [v_{cap}, v_0]$. This results in the following expression:

$$a_a(v) = \begin{cases} 0 & 0 \leq v < v_{cap} \\ -a_0 & v_{cap} \leq v < v_0 \\ -\kappa_a v & v_0 \leq v < v_{t,a} \\ -\kappa_a v_{t,a} \left( \frac{v_{\max,a} - v}{v_{\max,a} - v_{t,a}} \right) & v_{t,a} \leq v < v_{\max,a} \\ 0 & v \geq v_{\max,a} \end{cases} \tag{S.1}$$

Note that the assumption $a_a(v) = 0$ for $v < v_{cap}$ limits $v_a(t, v_b)$ to velocities greater than or equal to $v_{cap}$ (assuming that $v_b \geq v_{cap}$). While it is possible that some of the labeled magnetization may have reached a lower velocity, any potential error due to this assumption will be extremely small assuming typical values of $v_c$. For example, with $v_c = 2$ cm/s and a sinc saturation function, the value of $\text{sinc}(v_{cap}/v_c)$ is within 0.4% of sinc(0).

The expression for venous acceleration $a_v(v)$ has the same form but assumes that $v_0 = v_{cap}$ (so there are only 4 zones) and replaces the arterial transition velocity $v_{t,a}$ with the venous transition velocity $v_{t,v}$, the maximum arterial velocity $v_{\max,a}$ with the maximum venous velocity $v_{\max,v}$, and the negative of the arterial scaling parameter $-\kappa_a$ with the venous scaling parameter $\kappa_v$. Example waveforms are shown in Figure 3a with parameters $v_{cap} = 0.1$ cm/s, $\kappa_a = 5.04$ s$^{-1}$, $a_0 = 5.21$ cm/s$^2$, $v_0 = 0.42$ cm/s, $\kappa_v = 2.16$ s$^{-1}$, $v_{t,a} = v_{t,v} = 14$ cm/s, $v_{\max,a} = 30$ cm/s, and $v_{\max,v} = 23$ cm/s. The approach used to obtain these parameters is described in the SI Methods section.

##### S.1.2 Macrovascular outflow in the Local Model

In addition to the inflow of arterial blood described in Section 2.14, we must also account for macrovascular outflow in the local model, since an entering vessel may ultimately deliver blood to another voxel, either directly or through one of its branches. Note that we are still assuming a single compartment model, such that arterial blood that is ultimately delivered to a voxel decays at the blood relaxation rate [7]. We consider three cases:

1. The entering vessel and all its branches deliver blood to tissue in the voxel.

2. The entering vessel exits the voxel before branching and does not deliver any blood to tissue in the voxel.

3. The entering vessel branches within the voxel with some child vessels delivering blood to tissue in the voxel and others exiting the voxel. For each entering vessel we consider only one level of branching within the voxel.

For each entering vessel that has at least one associated vessel (either the same vessel or a child vessel) that exits the voxel, we denote the number of associated vessels (both exiting and non-exiting branches) for the $i$th vessel as $B_i(\tilde{\mathbf{r}})$, where $\tilde{\mathbf{r}}$ is the position-dimension vector defined in Section 2.14 and $B_i(\tilde{\mathbf{r}}) = 1$ for case 2 and $B_i(\tilde{\mathbf{r}}) \geq 1$ indicates the number of child vessels for case 3. For case 1, we set $B_i(\tilde{\mathbf{r}}) = 0$ since we do not need to track the outflow of any child vessels. For each exiting arterial vessel we denote the outflow velocity as $v_{o,i,j}(\tilde{\mathbf{r}}) \leq v_{b,i}(\tilde{\mathbf{r}})$ with exit index $j \in [1, \cdots, B_i(\tilde{\mathbf{r}})]$ and the fraction of the incoming flow $Q_0(\tilde{\mathbf{r}})f_i(\tilde{\mathbf{r}})$ that exits through the $j$th vessel as $g_{i,j}(\tilde{\mathbf{r}})$. If the vessel does not exit, then $g_{i,j}(\tilde{\mathbf{r}}) = 0$ and we can set $v_{o,i,j}(\tilde{\mathbf{r}}) = v_{b,i}(\tilde{\mathbf{r}})$ without loss of generality. Note that $\sum_{j=1}^{B_i(\tilde{\mathbf{r}})} g_{i,j}(\tilde{\mathbf{r}}) \leq 1$, with equality only if all child vessels exit. For a vessel that exits, the transit time from entry to exit can be expressed as

$$\Delta t_{v,i,j}(\tilde{\mathbf{r}}) = \int_{v_{b,i}(\tilde{\mathbf{r}})}^{v_{o,i,j}(\tilde{\mathbf{r}})} \frac{dv}{a_a(v)} \tag{S.2}$$

With the above definitions, the labeled arterial outflow from each voxel may be written as the weighted sum of time delayed arterial inflow functions

$$c_{a,0,\text{out}}(\tilde{\mathbf{r}}, t) = \sum_{i=1}^{N(\tilde{\mathbf{r}})} \sum_{j=1}^{B_i(\tilde{\mathbf{r}})} g_{i,j}(\tilde{\mathbf{r}}) \cdot c_{a,0,i}(\tilde{\mathbf{r}}, t - \Delta t_{v,i,j}(\tilde{\mathbf{r}})) \tag{S.3}$$

The overall arterial delivery function for the local model is the difference of the inflow and outflow terms

$$c_{a,0}(\tilde{\mathbf{r}}, t) = c_{a,0,\text{in}}(\tilde{\mathbf{r}}, t) - c_{a,0,\text{out}}(\tilde{\mathbf{r}}, t) \tag{S.4}$$

and the normalized cumulative signal is defined as

$$C_{a,0}(\tilde{\mathbf{r}}, t) = \frac{1}{\tau Q_0(\tilde{\mathbf{r}})} \int_0^t c_{a,0}(\tilde{\mathbf{r}}, u) du \tag{S.5}$$

To demonstrate the different cases, we consider three examples that are shown in Figure S6 for VSS with sinc saturation and in Figure S7 for VSS with windowed cosine saturation ($v_c$ = 2 cm/s for all examples). The case examples are as follows:

- Case 1 (panel a, inflow with two vessels): a voxel with $N(\tilde{\mathbf{r}}) = 2$ entering vessels with boundary velocities $v_{b,1}(\tilde{\mathbf{r}}) = 0.4$ cm/s and $v_{b,2}(\tilde{\mathbf{r}}) = 1.5$ cm/s and equal flow fractions $f_1(\tilde{\mathbf{r}}) = f_2(\tilde{\mathbf{r}}) = 0.5$ and no outflow from either vessel ($B_1(\tilde{\mathbf{r}}) = B_2(\tilde{\mathbf{r}}) = 0$). In this case the overall delivery function (solid blue) is the average of two delivery functions (gold and red) with a longer transit delay for $v_{b,1}(\tilde{\mathbf{r}}) = 0.4$ cm/s (red).

- Case 2 (panel c, one vessel passing through): a voxel with $N(\tilde{\mathbf{r}}) = 1$ entering vessel with boundary velocity $v_{b,1}(\tilde{\mathbf{r}}) = 1.5$ cm/s and $B_1(\tilde{\mathbf{r}}) = 1$ child vessel with exit velocity $v_{o,1,1}(\tilde{\mathbf{r}}) = 0.4$ cm/s. We assume that all entering blood exits the voxel ($g_{1,1}(\tilde{\mathbf{r}}) = 1$). In this case the overall delivery function (solid blue) consists of the difference between the inflow delivery function (red) and the outflow function (gold).

- Case 3 (panel e, two child vessels with one exiting): a voxel with $N(\tilde{\mathbf{r}}) = 1$ entering vessel with boundary velocity $v_{b,1}(\tilde{\mathbf{r}}) = 1.5$ cm/s; the vessel branches into $B_1(\tilde{\mathbf{r}}) = 2$ child vessels with no outflow from the first child vessel ($g_{1,1}(\tilde{\mathbf{r}}) = 0$) and 50 percent of the entering blood exiting the second child vessel ($g_{1,2}(\tilde{\mathbf{r}}) = 0.5$) with exit velocity $v_{o,1,2}(\tilde{\mathbf{r}}) = 0.4$ cm/s. In this case the overall delivery function (solid blue) reflects the inflow of blood with the first child vessel (gold) and the inflow and outflow of blood with the second child vessel (red).

The righthand columns of Figures S6 and S7 show the sum of the normalized cumulative signal $C_{a,0}(\tilde{\mathbf{r}}, t)$ and the initial arterial volume term at $t = 0$, which is non-zero only for vessels that deliver blood to the capillary bed. For reference, the sum signal from a single entering vessel with $v_{b,1}(\tilde{\mathbf{r}}) = 1.5$ cm/s is shown by the red dashed lines. This signal approaches 1.0, indicating complete delivery. For case 1, the sum signal lags the single vessel reference signal due to the presence of a second vessel with longer transit delay. In case 2, the sum signal approaches zero since all blood that enters eventually exits the vessel. Note however, that there is still substantial signal at $t = \tau = 1.4$s, which would manifest as an intravascular artifact in the voxel. Finally, for case 3, the sum signal approaches half the value of the reference signal, since half of the entering blood exits through the second child vessel.

##### S.1.3 PLD requirement for outflow in local model

For arterioles that pass through a voxel, the requirement is that any blood that is not saturated by the VCM has time to exit the voxel. This leads to the requirement PLD $> \Delta t_{i,j}(\tilde{\mathbf{r}})$ where $\Delta t_{i,j}(\tilde{\mathbf{r}})$ denotes the time required to decelerate from $v_c$ to $v_{o,i,j}(\tilde{\mathbf{r}})$. Since the exit velocity $v_{o,i,j}(\tilde{\mathbf{r}})$ of an arteriole exiting one voxel is the entry velocity for an adjacent voxel (i.e. $v_{b,i}(\tilde{\mathbf{r}} + \Delta\tilde{\mathbf{r}})$), the condition stated in Eq. 33 in the main text will tend to also be sufficient for the distribution of anticipated exit velocities. Note that there may also be an initial arterial volume term in the voxel that requires time to decelerate from to $v_{b,i}(\tilde{\mathbf{r}})$ to $v_{o,i,j}(\tilde{\mathbf{r}})$ in order to exit the voxel. The relevant requirement is that this time be less than $\tau + \mathrm{PLD}$, which is almost certainly met for typical values of $\tau$ and a PLD that satisfies Eq. 33.

#### S.2 Supporting Methods

We adapted a previously described vascular model [8] to estimate the scaling parameters and transition velocities used in the models of $a_a(v)$ and $a_v(v)$. The number of capillaries was decreased by 33% to obtain a mean capillary velocity of 1 mm/s, consistent with the range reported in a recent study [9]. Otherwise the model parameters are identical to those stated in Table 1 of [8]. Velocities were computed assuming an overall "normal" baseline cerebral blood flow of $Q_0 = 12.5$ ml/s (equivalent to 750 ml/min or 62.5 ml/(100g-min) for a 1.2 kg brain), which yielded arterial velocity versus diameter values similar to those reported in [10]. The time spent in each branch level was calculated as segment length divided by velocity, such that $t_{n,a} = l_{n,a}/v_{n,a}$ and $t_{n,v} = l_{n,v}/v_{n,v}$ where $n = 1$ denotes the smallest arteriole or venule with the index increasing with vessel size and the additional subscripts indicating the arterial (a) and venous (v) trees. The cumulative times in each tree (excluding the capillaries) were then calculated as $T_{n,a} = \sum_{i=1}^{n} t_{n,a}$ and $T_{n,v} = \sum_{i=1}^{n} t_{n,v}$. Figure 3b in the main text shows plots of velocity versus cumulative time for the arterial (red circles and dotted line) and the venous (blue x's and dotted line) branches for velocities below a transition velocity of $v_{t,a} = v_{t,v} = 14$ cm/s. In the vascular model, this transition velocity corresponds to vessels with diameters in the range of 0.4 to 0.6 mm, which is the estimated diameter for the start of the transition away from the Murray's law regime as described above in Section S.1.1.

Exponential fits to the velocity versus cumulative time curves were performed for velocities below the transition velocity. As noted above in Section S.1.1, for the arterial velocities, a better fit to the vascular model data was obtained by limiting the exponential fit to $v \in [v_0, v_{t,a}]$ and using an additional constant acceleration parameter for the arteriolar segment closest to the capillary bed ($v \in [v_{cap}, v_0]$). The solid lines in Figure 3b in the main text show the predicted velocity versus time curves ($v_a(t, v_{cap})$ and $v_v(t, v_{cap})$) using the parameters and the acceleration model described in Section S.1.1. For $v \leq 14$ cm/s the model curves exhibit an exponential dependence indicative of behavior in the Murray's law regime. For $v > 14$ cm/s, the models exhibit a transition to the da Vinci's rule regime, where the maximum arterial ($v_{\max,a} = 30$ cm/s) and venous ($v_{\max,v} = 23$ cm/s) velocities in the acceleration models were chosen to be consistent with literature values [11, 6]. In addition to the normal baseline flow parameters, we derived parameters assuming a lower value of $Q_0 = 7.7$ ml/s (equivalent to 460 ml/min or 38.3 ml/(100g-min)), which is the value used in [8]. The resulting model parameters for the low flow case were: $v_{cap} = 0.06$ cm/s, $\kappa_a = 2.99\ s^{-1}$, $a_0 = 1.96$

cm/$s^2$, $v_0 = 0.25$ cm/s, $\kappa_v = 1.25\ s^{-1}$, $v_{\max,a} = 18.4$ cm/s, and $v_{\max,v} = 14.1$ cm/s. The transition velocities $v_{t,a}$ and $v_{t,v}$ were held fixed over the normal and low flow cases. SI Figure S8 shows the acceleration versus velocity curves and the velocity versus time curves for both the normal and and low flow cases.

### A SI Appendix

#### A.1 Expressions for $v_a(t, v_b)$ and $v_v(t, v_{cap})$

We first derive an expression for the velocity $v_a(t, v_b)$ at $t = 0$ of an arterial spin that will decelerate to $v_b$ by time $t$. Given an arterial acceleration function $a_a(v) = \frac{dv}{dt} < 0$ we can write $\frac{dv}{a_a(v)} = dt$. Integrating both sides leads to

$$\int_{v_a(t,v_b)}^{v_b} \frac{dv}{a_a(v)} = -\int_{v_b}^{v_a(t,v_b)} \frac{dv}{a_a(v)} = \int_0^t du = t \tag{A.1}$$

where $v_a(t, v_b) \geq v_b$ for $t \geq 0$. Defining the indefinite integral $F_a(v) = -\int \frac{dv}{a_a(v)}$, we have $v_a(t, v_b) = {F_a}^{-1}(t + F_a(v_b))$ where ${F_a}^{-1}(u)$ denotes the inverse function of $F_a(v)$. As an example, when $a_a(v) = -\kappa_a v$ is valid over the velocity range of interest, $F_a(v) = \frac{1}{\kappa_a} \log v$ and ${F_a}^{-1}(u) = e^{\kappa_a u}$, so that $v_a(t, v_b) = v_b e^{\kappa_a t}$.

As noted in Section 2.4, when using expressions such as $v_a(t - \tau, v_b)$, the argument $t - \tau$ is negative over the range $[0, \tau)$. The solution forms derived in the previous paragraph are still valid when the argument is negative and can be used to evaluate $v_a(t - \tau, v_b)$. For example, with $a_a(v) = -\kappa_a v$ over the velocity range of interest, $v_a(t - \tau, v_b) = v_b e^{\kappa_a(t-\tau)}$, such that $v_a(t - \tau, v_b) < v_b$ for $t < \tau$.

For the specific model presented in Eq. S.1, $a_a(v) = 0$ for $0 \leq v < v_{cap}$ and $v \geq v_{\max,a}$, and therefore $F_a(v)$ is not defined in these regions. In practice, this is not an issue for $0 \leq v < v_{cap}$, since for computational purposes we can let $a_a(v) = -a_0$ in this region and then set $v_a(t, v_b) = v_{cap}$ when $v_a(t, v_b) < v_{cap}$. For $v_{t,a} \leq v < v_{\max,a}$, the form of $a_a(v)$ in Eq. S.1 leads to an exponential approach to $v_{\max,a}$ so that the case where $v \geq v_{\max,a}$ does not arise in the computation.

Next we derive the expression for the velocity $v_v(t, v_{cap})$ at time $t$ of a venous spin that starts with an initial velocity of $v_{cap}$ at $t = 0$. Given a venous acceleration function $a_v(v) > 0$, we have $\int_{v_{cap}}^{v_v(t,v_{cap})} \frac{dv}{a_v(v)} = t$. We can write $v_v(t, v_{cap}) = {F_v}^{-1}(t + F_v(v_{cap}))$ where the indefinite integral $F_v(v) = \int \frac{dv}{a_v(v)}$ and ${F_v}^{-1}(u)$ denotes the inverse function. As an example, if $a_v(v) = \kappa_v v$ is valid over the velocity range, then $F_v(v) = \frac{1}{\kappa_v} \log v$, ${F_a}^{-1}(u) = e^{\kappa_v u}$, and $v_v(t, v_{cap}) = v_{cap} e^{\kappa_v t}$.

#### A.2 Conditions for Ideal Bolus Width

As defined in Eq. 26, the effective bolus width is

$$\begin{aligned}\tau_{\text{eff}} &= \int_0^{\text{TI}} (1 - s_{\text{ref},E}(v_a(t, v_b))) \cdot s_{\text{ref}}(v_a(t-\tau, v_b))dt \\ &= \int_0^{\text{TI}} s_{\text{ref}}(v_a(t-\tau, v_b))dt - \int_0^{\text{TI}} s_{\text{ref},E}(v_a(t, v_b)) \cdot s_{\text{ref}}(v_a(t-\tau, v_b))dt \qquad \text{(A.2)}\end{aligned}$$

The first integral can be written as

$$\begin{aligned}\int_0^{\text{TI}} s_{\text{ref}}(v_a(t-\tau, v_b))dt &= \int_0^{\tau} s_{\text{ref}}(v_a(t-\tau, v_b))dt + \int_{\tau}^{\text{TI}} s_{\text{ref}}(v_a(t-\tau, v_b))dt \\ &= \tau + \underbrace{\int_0^{\tau} \Delta s_{\text{ref}}(v_a(t-\tau, v_b))dt}_{e_1} + \underbrace{\int_0^{\text{PLD}} s_{\text{ref}}(v_a(t, v_b))dt}_{A} \qquad \text{(A.3)}\end{aligned}$$

where PLD = TI $-\ \tau$ and $\Delta s_{\text{ref}}(v) = s_{\text{ref}}(v) - 1.0$ represents the deviation of $s_{\text{ref}}(v)$ from 1.0, which is its ideal value for $v \leq v_c$. Similarly, the second integral can be written as

$$\begin{aligned}-\int_0^{\text{TI}} s_{\text{ref},E}(v_a(t, v_b)) \cdot s_{\text{ref}}(v_a(t-\tau, v_b))dt &= \underbrace{-\int_0^{\tau} s_{\text{ref},E}(v_a(t, v_b))dt}_{B} \\ &\quad \underbrace{-\int_0^{\tau} s_{\text{ref},E}(v_a(t, v_b))\Delta s_{\text{ref}}(v_a(t-\tau, v_b))dt}_{e_2} \\ &\quad \underbrace{-\int_0^{\text{PLD}} s_{\text{ref},E}(v_a(t+\tau, v_b)) \cdot s_{\text{ref}}(v_a(t, v_b))dt}_{C} \qquad \text{(A.4)}\end{aligned}$$

Summing the expressions yields $\tau_{\text{eff}} = \tau + A - B - C + e_1 - e_2$. Sufficient conditions for $\tau_{\text{eff}} = \tau$ are that $A = B$ and $C = e_1 = e_2 = 0$.

We consider the typical VSS case where the LCM and VCM use the same saturation pulse sequence modules, so that $s_{\text{ref},E}(v) = s_{\text{ref}}(v)$ and $p_{\text{ref}}(v) = 1 - s_{\text{ref}}(v)$. With this assumption, a sufficient condition for $C = 0$ is that $s_{\text{ref}}(v_a(t, v_b))$ and $s_{\text{ref}}(v_a(t+\tau, v_b))$ do not overlap over the interval $0 \leq t \leq$ PLD. A straightforward way to satisfy this condition is to require that $s_{\text{ref}}(v_a(t, v_b)) = 0$ for $t \geq \tau$, since this leads to $s_{\text{ref}}(v_a(t+\tau, v_b)) = 0$ for $t \geq 0$. This is equivalent to requiring that $s_{\text{ref}}(v) = 0$ for velocities above a cutoff velocity $v_c$, where $v_c \leq v_\tau$ is less than $v_\tau = v_a(\tau, v_b)$. This is **Condition W1**. If Condition W1 is met, then we also have $A = B$ as long as PLD $\geq \tau$ or TI $\geq 2\tau$. Furthermore, Condition W1 implies the requirement $\tau \geq \Delta t$, which is necessary in order to achieve $v_c = v_a(\Delta t, v_b) \leq v_a(\tau, v_b)$.

For typical values of $\tau$ on the order of one second or more, $v_\tau$ will approach the maximum arterial velocities (e.g. 30 cm/s as shown in Figure 3 in the main text), which is much greater than recommended values for $v_c$ (e.g. 2 cm/s). Thus for typical values of $v_c$, the transit time $\Delta t$ required for arterial spins to decelerate from $v_c$ to $v_b$ is less than $\tau$. With Condition W1 we have $s_{\text{ref}}(v) = 0$ for $v \geq v_c$, so that the integrand for part $B$ is non-zero only over the interval $0 \leq t \leq \Delta t$. As a result, we will have $A = B$ as long as PLD $\geq \Delta t$. This is **Condition W2**.

We now consider the error terms $e_1$ and $e_2$. Making use of the relations $v_a(t-\tau, v_b) = v_a(t-\tau+\Delta t_b, v_{cap})$ and $\Delta s_{\text{ref}}(v) = -p_{\text{ref}}(v)$ (assuming VSS), we can write the error term $e_1$ as

$$\begin{aligned} e_1 &= \int_0^{\tau} \Delta s_{\text{ref}}(v_a(t-\tau+\Delta t_b, v_{cap}))dt \\ &= \int_{-\tau+\Delta t_b}^{\Delta t_b} \Delta s_{\text{ref}}(v_a(t, v_{cap}))dt \\ &= \int_{-\tau+\Delta t_b}^{0} \Delta s_{\text{ref}}(v_a(t, v_{cap}))dt - \int_0^{\Delta t_b} p_{\text{ref}}(v_a(t, v_{cap}))dt \\ &= (\tau - \Delta t_b) \cdot \Delta s_{\text{ref}}(v_{cap}) - V_a(0)/(Q_0 \cdot P_a) \\ &\approx -V_a(\tau)/(Q_0 \cdot P_a \cdot S_0) \end{aligned} \tag{A.5}$$

where $V_a(0)$ and $V_a(\tau)$ are arterial volume terms and $\Delta t_b$ is the transit time from $v_b$ to $v_{cap}$ (see definitions and equations in Section 2.5). The final approximation reflects the fact that $\Delta s_{\text{ref}}(v_{cap}) \approx 0$ for most implementations. For the global model $e_1 \approx 0$ since by definition $V_a(0) = 0$ when $v_b = v_{cap}$. In the local model, the term $e_1$ approximately cancels out the component $V_a(\tau)/(Q_0 \cdot P_a \cdot S_0)$ of the effective bolus width associated with the arterial volume term (see Section 2.14 and Figure 4 in the main text for further discussion).

For the term $e_2$, we note that $s_{\text{ref}}(v_a(t, v_b)) = 0$ for $t > \Delta t$ when Condition W1 holds. Furthermore, $\Delta s_{\text{ref}}(v_a(t-\tau, v_b)) \approx 0$ for $t < \tau - \Delta t_b$, since $\Delta s_{\text{ref}}(v_{cap}) \approx 0$. Thus, for $\tau > \Delta t + \Delta t_b$, the product $s_{\text{ref}}(v_a(t, v_b)) \cdot \Delta s_{\text{ref}}(v_a(t-\tau, v_b)) \approx 0$ over the interval $[0, \tau]$ and thus $e_2 \approx 0$. For typical acquisitions where $\tau >$ PLD and assuming that Condition W2 holds, the condition $\tau > \Delta t + \Delta t_b$ is satisfied for the global model (since $\Delta t_b = 0$ when $v_b = v_{cap}$), and will also be satisfied for the local model as long as $\tau >$ PLD $+ \Delta t_b$.

To summarize, when conditions W1 and W2 are met for VSS with $s_{\text{ref},E}(v) = s_{\text{ref}}(v)$, then we have $A = B$, $C \approx e_2 \approx 0$, and $e_1$ is either close to zero in the global model or cancels out the arterial volume term in the local model, resulting in $\tau_{\text{eff}} = \tau$.

#### A.3 Bolus Width Error when Condition W2 is not met

We assume that (i) $s_{\text{ref},E}(v) = s_{\text{ref}}(v)$, (ii) condition W1 is satisfied so that $C = 0$ and $\tau \geq \Delta t$, and furthermore (iii) $\tau > \Delta t + \Delta t_b$ so that $e_2 \approx 0$. We also assume that that Condition W2 is not satisfied so that PLD $< \Delta t$. Adapting the expressions from section A.2, we can write the bolus width error $\Delta\tau_{\text{eff}} = \tau_{\text{eff}} - \tau = A - B$ as

$$\begin{aligned} \Delta\tau_{\text{eff}} &= \int_0^{\text{PLD}} s_{\text{ref}}(v_a(t, v_b))dt - \int_0^{\Delta t} s_{\text{ref}}(v_a(t, v_b))dt \\ &= -\int_{\text{PLD}}^{\Delta t} s_{\text{ref}}(v_a(t, v_b))dt \end{aligned} \tag{A.6}$$

#### A.4 Mismatch case

Now we consider what happens when $s_{\text{ref},E}(v) \neq s_{\text{ref}}(v)$. We assume that the $s_{\text{ref},E}(v)$ and $s_{\text{ref}}(v)$ are zero above their respective cutoff velocities of $v_{l,\text{eff}}$ and $v_c$, where both cutoff velocities are less than $v_\tau$ such that Condition W1 is met (so $C = 0$), and we have used the effective labeling cutoff velocity $v_{l,\text{eff}}$ instead of the nominal cutoff velocity $v_l$. Since there are two cutoff velocities, it is no longer sufficient to define a single transit delay, and therefore we define the LCM and VCM transit delays as the times $\Delta t_l$ and $\Delta t_c$ required for blood to decelerate from $v_{l,\text{eff}}$ or $v_c$, respectively, to $v_b$. We also define $\Delta t_{\max} = \max(\Delta t_l, \Delta t_c)$.

Following the reasoning from Section A.2, we have $e_2 \approx 0$ when $\tau > \Delta t_l + \Delta t_b$. We can also show that

$$e_1 \approx -V_a(\tau)/(Q_0 \cdot P_a \cdot S_0) + \underbrace{\int_0^{\Delta t_b} (s_{\text{ref}}(v_a(t, v_{cap})) - s_{\text{ref},E}(v_a(t, v_{cap})))dt}_{\Delta e_1} \tag{A.7}$$

is equal to sum of the volume-related term previously derived in Eq. A.5 and a correction term $\Delta e_1$. This correction term is zero for the global model and tends to be close to zero for the local model when the saturation functions are similar in shape for low velocities in the range $[v_{cap}, v_b]$.

Focusing on the remaining components, the effective bolus width error can be written as

$$\Delta\tau_{\text{eff}} = A - B = \int_0^{\text{PLD}} s_{\text{ref}}(v_a(t, v_b))dt - \int_0^{\tau} s_{\text{ref},E}(v_a(t, v_b))dt \tag{A.8}$$

Under the condition that PLD $\geq \Delta t_{\max}$, we can write

$$\begin{aligned} \Delta\tau_{\text{eff}} &= \int_0^{\Delta t_{\max}} \left(s_{\text{ref}}(v_a(t, v_b)) - s_{\text{ref},E}(v_a(t, v_b))\right) dt \\ &= \int_{v_b}^{\max(v_{l,\text{eff}}, v_c)} \frac{(s_{\text{ref},E}(v) - s_{\text{ref}}(v))}{a_a(v)} dv \end{aligned} \tag{A.9}$$

Examining the form of Eq. A.9 and noting that $a_a(v) < 0$, we can see that if $s_{\text{ref},E}(v)$ has a broader response (i.e. $v_{l,\text{eff}} > v_c$), then $\Delta\tau_{\text{eff}} < 0$ will be negative. This corresponds to the VCM function prematurely saturating labeled spins that would have otherwise been delivered. On the other hand, if $s_{\text{ref}}(v)$ has a broader response (i.e. $v_c > v_{l,\text{eff}}$) then $\Delta\tau_{\text{eff}} > 0$ will be positive. In this case the higher cutoff velocity of the VCM allows for the delivery of excess labeled blood. Note that Eq. A.9 assumes that $s_{\text{ref},E}(v)$ and $s_{\text{ref}}(v)$ have finite support with hard cutoff velocities. In practice, Eq. A.9 still serves as a good approximation for saturation functions (e.g. sinc saturation and VSI effective saturation function) that do not have a hard cutoff velocity but oscillate about zero above their nominal cutoff frequencies.

#### A.5 Local model arterial delivery and volume terms

Adopting the notation of Section 2.14, we consider the case of a single arteriole ($N(\tilde{\mathbf{r}}) = 1$) delivering blood to the capillary beds in a voxel with input flow $Q_{0,1}(\tilde{\mathbf{r}})$. We can write the cumulative label delivered to the intra-voxel capillary beds ($v_b = v_{cap}$) over a time interval $[0, \text{TI}]$ as

$$\begin{aligned}
\int_0^{\text{TI}} c_{a,0}(\tilde{\mathbf{r}}, t, v_{cap})dt &= Q_{0,1}(\tilde{\mathbf{r}}) \cdot \int_0^{\text{TI}} p(v_a(t, v_{cap})) \cdot s(v_a(t-\tau, v_{cap}))dt \\
&= Q_{0,1}(\tilde{\mathbf{r}}) \cdot \int_0^{\Delta t_{b,1}} p(v_a(t, v_{cap})) \cdot s(v_a(t-\tau, v_{cap}))dt + \\
&\quad\; Q_{0,1}(\tilde{\mathbf{r}}) \cdot \int_{\Delta t_{b,1}}^{\text{TI}} p(v_a(t, v_{cap})) \cdot s(v_a(t-\tau, v_{cap}))dt \\
&= V_a(\tilde{\mathbf{r}}, \tau) + Q_{0,1}(\tilde{\mathbf{r}}) \cdot \int_0^{\text{TI}-\Delta t_{b,1}} p(v_a(t, v_{b,1}(\tilde{\mathbf{r}}))) \cdot s(v_a(t-\tau, v_{b,1}(\tilde{\mathbf{r}})))dt \\
&= V_a(\tilde{\mathbf{r}}, \tau) + \int_0^{\text{TI}-\Delta t_{b,1}} c_{a,0,1}(\tilde{\mathbf{r}}, t)dt \qquad \text{(A.10)}
\end{aligned}$$

where we introduce the notation $c_{a,0}(\tilde{\mathbf{r}}, t, v_{cap})$ in the first line to represent delivery to the capillary beds in the voxel. In addition, $v_{b,1}(\tilde{\mathbf{r}})$ denotes the boundary velocity of the feeding arteriole, $\Delta t_{b,1}$ is the transit delay from $v_{b,1}(\tilde{\mathbf{r}})$ to $v_{cap}$ and

$$V_a(\tilde{\mathbf{r}}, \tau) = Q_{0,1}(\tilde{\mathbf{r}}) \cdot \int_0^{\Delta t_{b,1}} p(v_a(u, v_{cap})) \cdot s(v_a(u-\tau, v_{cap}))du \qquad \text{(A.11)}$$

is the local version of the global arterial volume term defined in Eq. 9. Note that we have made use of the fact that $v_a(t + \Delta t_{b,1}, v_{cap}) = v_a(t, v_{b,1}(\tilde{\mathbf{r}}))$. Eq. A.10 states that the labeled blood delivered to the capillary bed in a voxel over an interval $[0, \text{TI}]$ can be expressed as the sum of the labeled blood delivered to the voxel over an interval $[0, \text{TI} - \Delta t_{b,1}]$ and the volume of labeled arterial blood that is created by the LCM at $t = 0$ and remains withins the voxel. This volume of blood is delivered to the capillary beds in the voxel over the interval $[0, \Delta t_{b,1}]$.

Note that if $\text{TI} = \tau + \Delta t$ where $\Delta t$ is the transit time from $v_c$ to $v_{cap}$, then Eq. A.10 states that the required inflow time for the local model is $\text{TI}' = \text{TI} - \Delta t_{b,1}$, which is equivalent to $\text{TI}' = \tau + \Delta t - \Delta t_{b,1} = \tau + \Delta t'$ where $\Delta t' = \Delta t - \Delta t_{b,1}$ is the transit delay from $v_c$ to $v_b$. This is also equivalent to stating that the required PLD for this local model example is $\text{PLD}_{local} = \text{PLD}_{global} - \Delta t_{b,1}$.

#### A.6 Capillary and venous blood volume terms

*Capillary Blood Volume component.* The volume of labeled magnetization in the capillaries is modeled as $V_{cap}(0) = Q_0 \cdot t_{cap} \cdot p(v_{cap})$ where $t_{cap}$ is the mean capillary transit time. Since $p(v_{cap}) \approx 0$ for almost all VSASL implementations that are of interest, we will ignore the contribution of this term to minimize the complexity of the presentation.

*Venous Blood Volume component.* The volume of venous blood in the range $[v_{cap}, v_{\max,v}]$ is equal to $Q_0 \cdot \Delta t_{V,\max}$ where $v_{\max,v}$ denotes the maximum venous velocity, $\Delta t_{V,\max}$ is the time required for blood to accelerate from $v_{cap}$ to $v_{\max,v}$, and we assume that blood at $v_{\max,v}$ is already out of the region of interest so there is no need to consider any additional contribution from blood that stays at this maximum velocity. Weighting each time increment of blood according to the passband value at each increment's initial velocity, we obtain the following expression for the venous blood volume $V_v$ created by the LCM at $t = 0$:

$$V_v(0) = Q_0 \cdot \int_0^{\Delta t_{V,\max}} p(v_v(u, v_{cap}))du \tag{A.12}$$

where $v_v(u,v_{cap}) \geq v_{cap}$ denotes the velocity at time $u$ of venous blood that started at $v_{cap}$ (see also Appendix section A.1). At $t = \tau$, the VCM will saturate each time increment of labeled blood volume according to the velocity that it has accelerated to. This can be expressed as

$$V_v(\tau) = Q_0 \cdot \int_0^{\Delta t_{V,\max}} p(v_v(u, v_{cap}))s(v_v(u + \tau, v_{cap}))du \tag{A.13}$$

We now examine the terms in Eq. A.13 for the VSS case where $p(v) = 1 - s(v)$ (assuming $S_0 = 1$ to simplify the presentation). Assuming that Condition W1 holds with some cutoff velocity $v_c$, we then have $s(v_v(u + \tau, v_{cap})) = 0$ for $u + \tau \geq \Delta t_{V,c}$ where $\Delta t_{V,c}$ denotes the time for venous blood to accelerate from $v_{cap}$ to $v_c$. If $\tau > \Delta t_{V,c}$, then $s(v_v(u + \tau, v_{cap})) = 0$ starting at a negative value $u = \Delta t_{V,c} - \tau$, resulting in $V_v(\tau) = 0$ under the assumption of hard cutoff velocities and $V_v(\tau) \approx 0$ for profiles that oscillate about zero above their nominal cutoff velocities. If $\tau \leq \Delta t_{V,c}$, then we further require that $p(v_v(u, v_{cap})) = 0$ over the interval $u \in [0, \Delta t_{V,c} - \tau]$. This is equivalent to requiring that $s(v) = 1$ for $v \leq v_v(\Delta t_{V,c} - \tau, v_{cap})$. Based on the estimates of $v_v(t, v_{cap})$ shown in Figure S8b and assuming $v_c = 2$ cm/s and $\tau = 1.4$ s, we will typically be in a regime where either (1) $\tau > \Delta t_{V,c}$ or (2) $v_v(\Delta t_{V,c} - \tau, v_{cap})$ is not much greater than $v_{cap}$ so that the $s(v) \approx s(v_{cap}) = 1$ for $v \leq v_v(\Delta t_{V,c} - \tau, v_{cap})$ is a reasonable approximation. In both regimes, we therefore have $V_v(\tau) \approx 0$. For the VSI mismatch case where $v_l = v_c$ and $v_{l,\text{eff}} > v_c$, then the intervals over which $p(v_v(u, v_{cap}))$ and $s(v_v(u + \tau, v_{cap}))$ are zero will overlap, and again $V_v(\tau) \approx 0$. Overall, we expect that $V_v(\tau) \approx 0$ for most cases of interest, but there may be edge cases (e.g. $v_{l,\text{eff}} < v_c$ and very small values of $\tau$) where it is helpful to use Eq. A.13 to evaluate the magnitude of the term.

#### A.7 Potential extensions to definition of the passband function

Following the prior work [12, 13], we have assumed laminar flow profiles to obtain passband functions of the form

$$p(\bar{v}) = \frac{1}{2\bar{v}} \int_0^{2\bar{v}} p_0(v)dv \tag{A.14}$$

where $p_0(v)$ denotes the passband response prior to laminar flow integration. The same approach was also adopted for the form of the saturation functions $s(v)$.

In our model we use terms of the form $p(v_a(t, v_b))$, where $v_a(t, v_b)$ denotes the mean velocity of blood that will decelerate to $v_b$ at time $t$. Thus, we may also write

$$p(v_a(t, v_b)) = \frac{1}{2v_a(t, v_b)} \int_0^{2v_a(t,v_b)} p_0(v)dv \tag{A.15}$$

To account for the varying probability that blood at different radial positions in the vessel will decelerate to $v_b$ at time $t$, we can introduce the following modification

$$p(v_a(t, v_b)) = \int_0^{2v_a(t,v_b)} \int_{t-\epsilon/2}^{t+\epsilon/2} p_0(v) \cdot f(v, t'; v_a(t, v_b), v_b)dt'dv \tag{A.16}$$

where $\epsilon$ is on the order of the simulation time increment (e.g. 1 ms) and $f(v, t'; v_a(t, v_b), v_b)dt'dv$ denotes the probability that blood with velocity $v \in [0, 2v_a(t, v_b)]$ in vessels with mean velocity $v_a(t, v_b)$ will decelerate to $v_b$ at time $t'$. The choice $f(v, t'; v_a(t, v_b), v_b) = \frac{1}{2v_a(t,v_b)}\delta(t' - t)$ yields Eq. A.15, where $\delta(\cdot)$ denotes the Dirac delta function. For specified values of $t$ and $v_b$, the normalization of the probability density function is

$$\int_0^{2v_a(t,v_b)} \int_0^{\infty} f(v, t'; v_a(t, v_b), v_b)dt'dv = 1, \tag{A.17}$$

Next to allow for the possibility that blood from various points along the arterial tree may reach $v_b$ in time $t$, we further extend the definition as follows

$$p(v_a(t, v_b)) = \int_{-t}^{\infty} \chi(\eta) \int_0^{2v_a(t+\eta,v_b)} \int_{t+\eta-\epsilon/2}^{t+\eta+\epsilon/2} p_0(v) \cdot f(v, t'; v_a(t + \eta, v_b), v_b)dt'dvd\eta \tag{A.18}$$

where $\chi(\eta)d\eta$ denotes the fraction of blood flow that reaches the boundary $v_b$ at time $t$ from vessels with mean velocity $v_a(t+\eta, v_b)$ where $\eta \in [-t, \infty]$ and $\int_{-t}^{\infty} \chi(\eta)d\eta = 1$. Note that if $\chi(\eta) = \delta(\eta)$ is a Dirac delta function and $f(v, t'; v_a(t, v_b), v_b) = \frac{1}{2v_a(t,v_b)}\delta(t'-t)$, then the passband function in Eq. A.18 is equivalent to the passband function under the assumption of laminar flow as described in Eq. A.15.

Eq. A.18 represents one potential approach for going beyond the laminar flow profile assumption to derive passband functions that more fully reflect the complexity of microvascular flow. However, that same complexity is likely to make it challenging to obtain useful representations of the functions $f(v, t'; v_a(t, v_b), v_b)$ and $\chi(\eta)$.

#### A.8 Approximation for FTVSI passband function

As demonstrated by the examples for $v_l = 1.2$ and 2 cm/s in SI Figure S10(a,b), a reasonable empirical approximation for the FTVSI passband function prior to integration for laminar flow is

$$p_0(v) = \begin{cases} 1 - \cos\left(\frac{2\pi(v - kLv_l)}{4v_l}\right) & (kL-2)v_l \le v < (kL+2)v_l \\ 2 & \text{otherwise} \end{cases} \tag{A.19}$$

where $k$ spans the integers, the aliased components in the response occur at multiples of $Lv_l$, and $L$ is an integer that depends on the sampling of velocity k-space in the LCM.

To obtain the passband function assuming laminar flow with mean velocity $\bar{v}$, we take the average of $p_0(v)$ over an interval $[0, 2\bar{v}]$ to yield the following approximation:

$$p_{\text{lam}}(\bar{v}) = \begin{cases} 1 - \text{sinc}(\bar{v}/v_l) & 0 \le |\bar{v}| < v_l \\ 2 - \frac{(2k+1)v_l}{|\bar{v}|} & \frac{(kL+2)v_l}{2} \le |\bar{v}| < \frac{((k+1)L-2)v_l}{2} \\ 1 + \frac{(L-4)(k+1)v_l}{2|\bar{v}|} + \frac{\sin\left(\frac{\pi\left(|\bar{v}| - ((k+1)L-2)v_l/2\right)}{v_l}\right)}{\pi|\bar{v}|/v_l} & \frac{((k+1)L-2)v_l}{2} \le |\bar{v}| < \frac{((k+1)L+2)v_l}{2} \end{cases} \tag{A.20}$$

where $k$ spans the non-negative integers. The laminar passband functions obtained from Bloch equation simulations for $v_l = 1.2$ and 2 cm/s and the approximations are shown in SI Figure S10(c), with the corresponding approximation errors shown in panel (d). Because of the averaging process, the errors in the laminar flow approximation (panel (d)) are about one order of magnitude smaller than those for the plug flow approximation (panel (b)). It can be shown that the steady state value of $p_{\text{lam}}(\bar{v})$ is $P_a = 2 - 4/L$. Note that in keeping with the convention stated in Section 2, we have used the $\bar{v}$ notation to clarify the derivation, but will revert back to using $v = \bar{v}$ when referring to the laminar response.

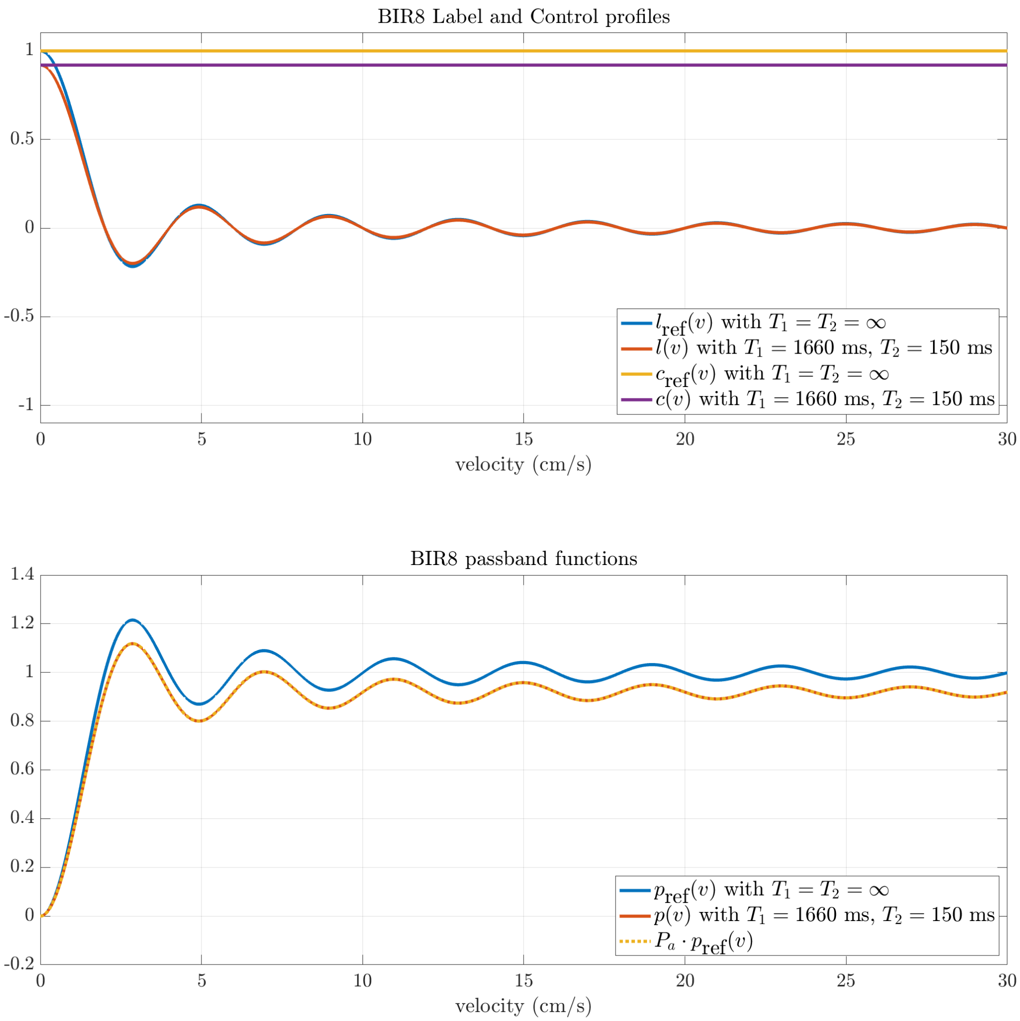


Figure S1: (a) Bloch simulated BIR8 label and control profiles with either $T_1 = T_2 = \infty$ for reference profiles ($l_{\text{ref}}(v)$ and $c_{\text{ref}}(v)$) and $T_1 = 1660$ ms and $T_2 = 150$ ms for $l(v)$ and $c(v)$. As compared to the reference profiles, $l(v)$ and $c(v)$ are both scaled by a factor of $P_a = 0.92$ (see Methods for additional simulation parameters). (b) Passband functions $p_{\text{ref}}(v) = c_{\text{ref}}(v) - l_{\text{ref}}(v)$ and $p(v) = c(v) - l(v)$ corresponding to the label and control profiles shown in panel (a). Note that $p(v) = P_a \cdot p_{\text{ref}}(v)$ as shown by the matching of the red solid curve and yellow dotted curve.

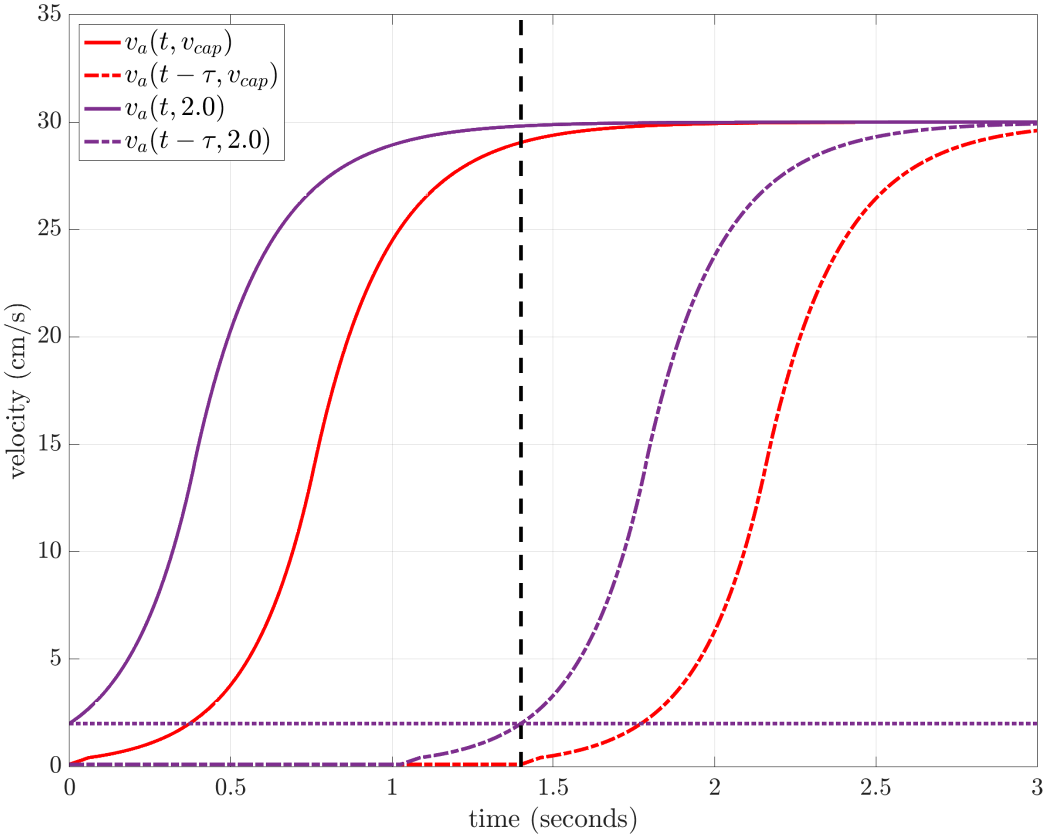


Figure S2: Plots of $v_a(t, v_b)$ (solid) and $v_a(t-\tau, v_b)$ (dash) for $v_b = v_{cap} = 0.1$ cm/s (red) and $v_b = 2.0$ cm/s (magenta) calculated using the arterial acceleration model parameters from Section S.1.1. The black vertical line indicates $t = \tau$. At $t = \tau$, we have $v_a(t-\tau, v_b) = v_a(0, v_b) = v_b$, and hence the intersection of the $v_a(t-\tau, v_b)$ dashed curves with the black vertical line occur at the points $(\tau, v_b)$. For $v_b = 2.0$ cm/s, this point is indicated by the intersection of the black vertical line with the horizontal magenta line. For times $t \leq \tau$, the argument $t - \tau$ is negative, resulting in $v_a(t-\tau, v_b)$ taking on values less than or equal to $v_b$.

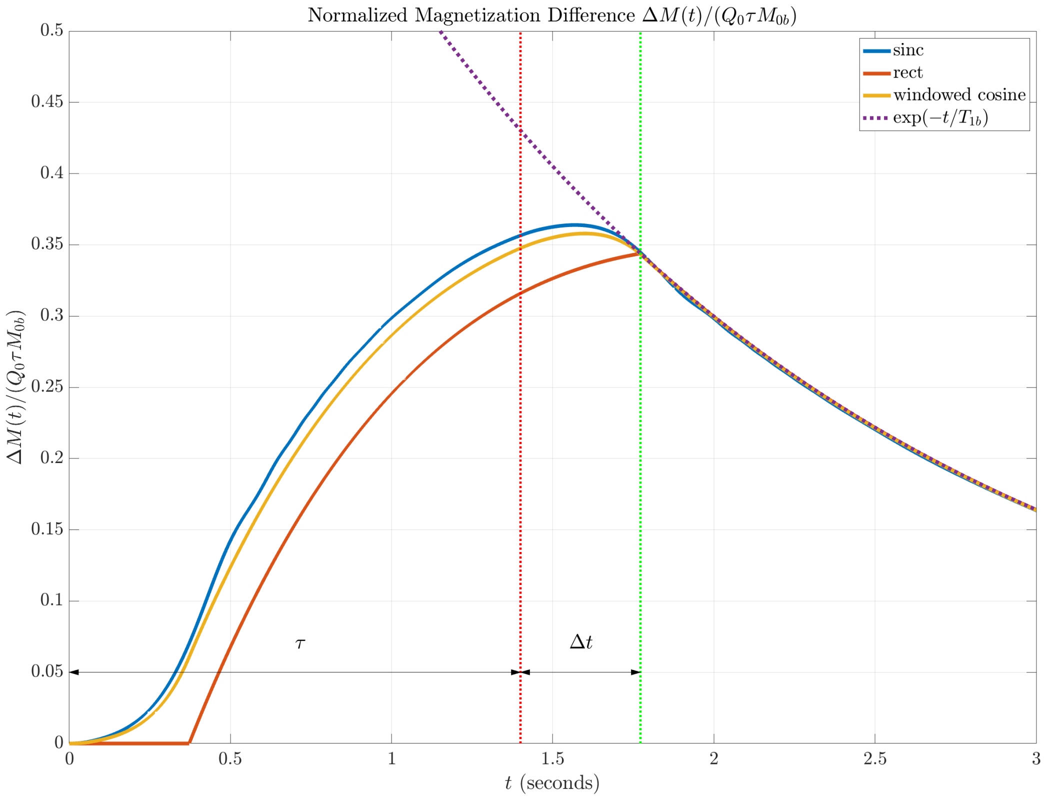


Figure S3: Plots of normalized magnetization difference $\widehat{\Delta M} = \Delta M(t)/(M_{0b} Q_0 \tau)$ for VSS with sinc (blue), rect (red), and windowed cosine (yellow) saturation functions and $P_a = S_0 = 1$. The dotted purple line shows the exponential decay curve $\exp(-t/T_{1b})$ with $T_{1b} = 1660$ ms. Note that the $\widehat{\Delta M}$ curves can be written as the product $\widehat{\Delta M} = \exp(-t/T_{1b}) \cdot C_{a,0}(t)$ of the exponential decay curve and the normalized cumulative time integral $C_{a,0}(t)$ curves shown in Figure 1d. Since the $C_{a,0}(t)$ curves approach their steady-state value of 1.0 at $t = \tau + \Delta t$, the $\widehat{\Delta M}$ curves follow the exponential decay curve $\exp(-t/T_{1b})$ for $t \geq \tau + \Delta t$.

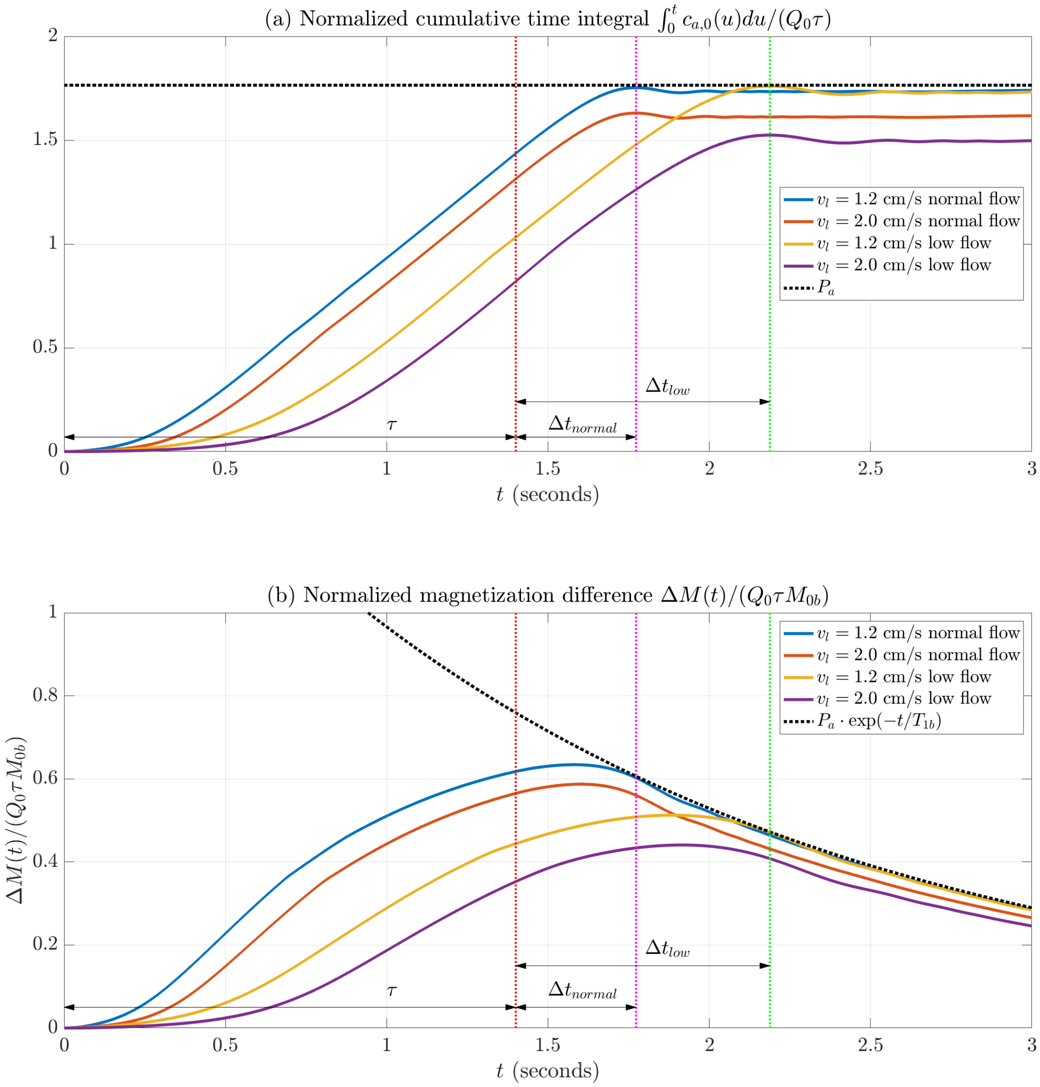


Figure S4: (a) Normalized cumulative time integrals $C_{a,0}(t)$ for VSI with $v_l = 1.2$ cm/s and $v_l = 2$ cm/s for both normal (red, blue) and low flow conditions (yellow, purple), with $S_0 = 1.0$ and $P_a = 1.7647$ (black dotted line). (b) Plots of normalized magnetization difference $\widehat{\Delta M} = \Delta M(t)/(M_{0b}Q_0\tau) = \exp(-t/T_{1b}) \cdot C_{a,0}(t)$ The dotted black line shows the curve $P_a \cdot \exp(-t/T_{1b})$ where $T_{1b} = 1660$ ms. Note that the $C_{a,0}(t)$ curves for $v_l = 1.2$ cm/s approach the steady-state value of $P_a$ at $t = \tau + \Delta t_{normal}$ and $t = \tau + \Delta t_{low}$ for normal and low flow conditions, respectively. As a result, the $\widehat{\Delta M}$ curves for $v_l = 1.2$ approximately follow the exponential decay curve $P_a \cdot \exp(-t/T_{1b})$ for $t \geq \tau + \Delta t_{normal}$ and $t \geq \tau + \Delta t_{low}$ for normal and low flow conditions, respectively. The lower relative amplitudes of both the $C_{a,0}(t)$ and $\widehat{\Delta M}$ curves for $v_l = 2.0$ cm/s reflect the effects of the mismatch discussed in Section 2.13.

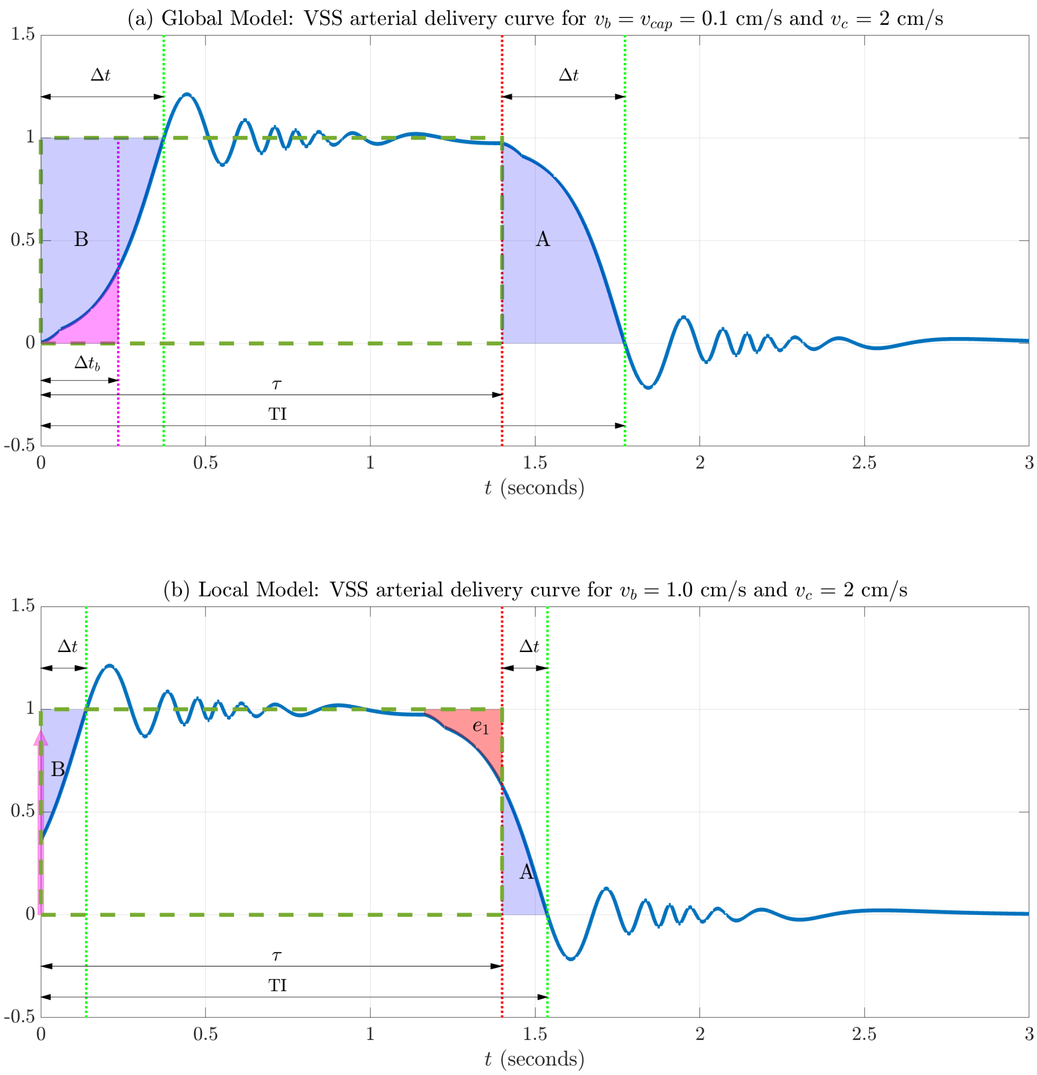


Figure S5: Arterial delivery components for the global and local models, assuming VSS with a sinc saturation function. (a) When considering delivery to the capillary beds, the transit delay $\Delta t$ is the time for blood to decelerate from the $v_c = 2$ cm/s to $v_{cap} = 0.1$ cm/s. As long as PLD $\geq \Delta t$, the leading and trailing edges are complementary and the areas A and B match. As a result, the integral of the arterial delivery function (solid blue curve) over the interval $[0, \mathrm{TI}]$ is equal to the area of the rectangle (dashed green line) and $\tau_{\mathrm{eff}} = \tau$. (b) In the local model, $\Delta t$ is the time required to decelerate to from $v_c = 2$ cm/s to $v_b = 1$ cm/s. As long as PLD $\geq \Delta t$, then areas A and B match. In addition, the error term $e_1$ is approximately matched by the labeled blood volume component (magenta arrow) injected at $t = 0$, so that the $\tau_{\mathrm{eff}} \approx \tau$. This volume component is equal to the magenta area in panel (a), where $\Delta t_b$ is the time needed to decelerate from $v_b$ to $v_{cap}$.

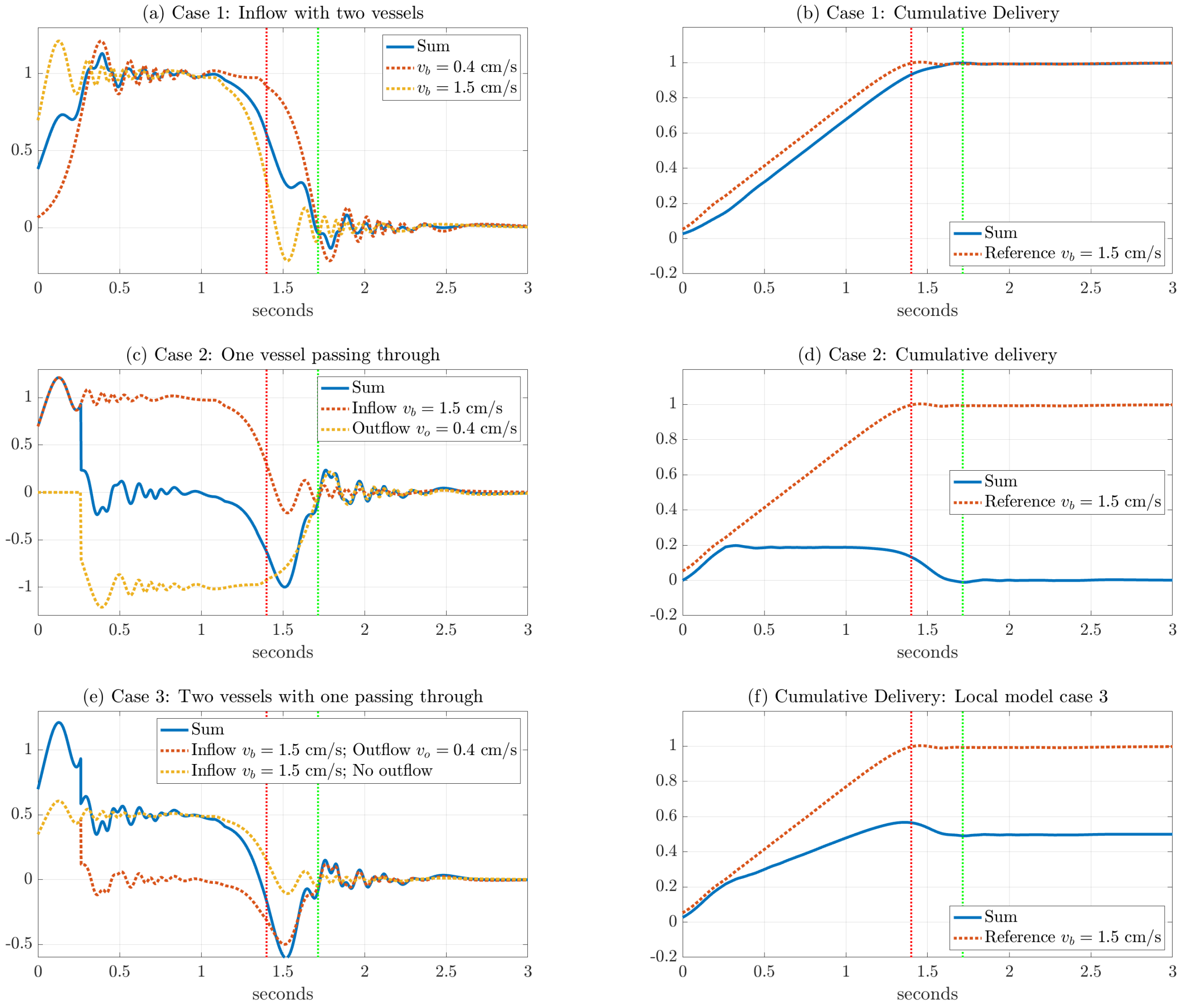


Figure S6: Arterial delivery functions (panels a, c, e) and cumulative delivery signals (panels b, d, f) for the local model case examples, assuming sinc VSS with $v_c = 2$ cm/s. Note that the arterial delivery functions are normalized by $Q_0(\tilde{\mathbf{r}})$, and the cumulative delivery signals include the effect of an initial arterial volume term for vessels that deliver blood to the capillary bed. Case 1 (panels a, b): two arterioles with different boundary velocities (0.4 cm/s and 1.5 cm/s) deliver blood to the capillary bed. Case 2 (panels c, d): an arteriole passes through the voxel with entry and exit velocities of 1.5 cm/s and 0.4 cm/s, respectively. Case 3 (panels e, f): arteriole that branches into two child arterioles, with one delivering blood to the capillary bed and the other exiting the voxel. See text for additional details.

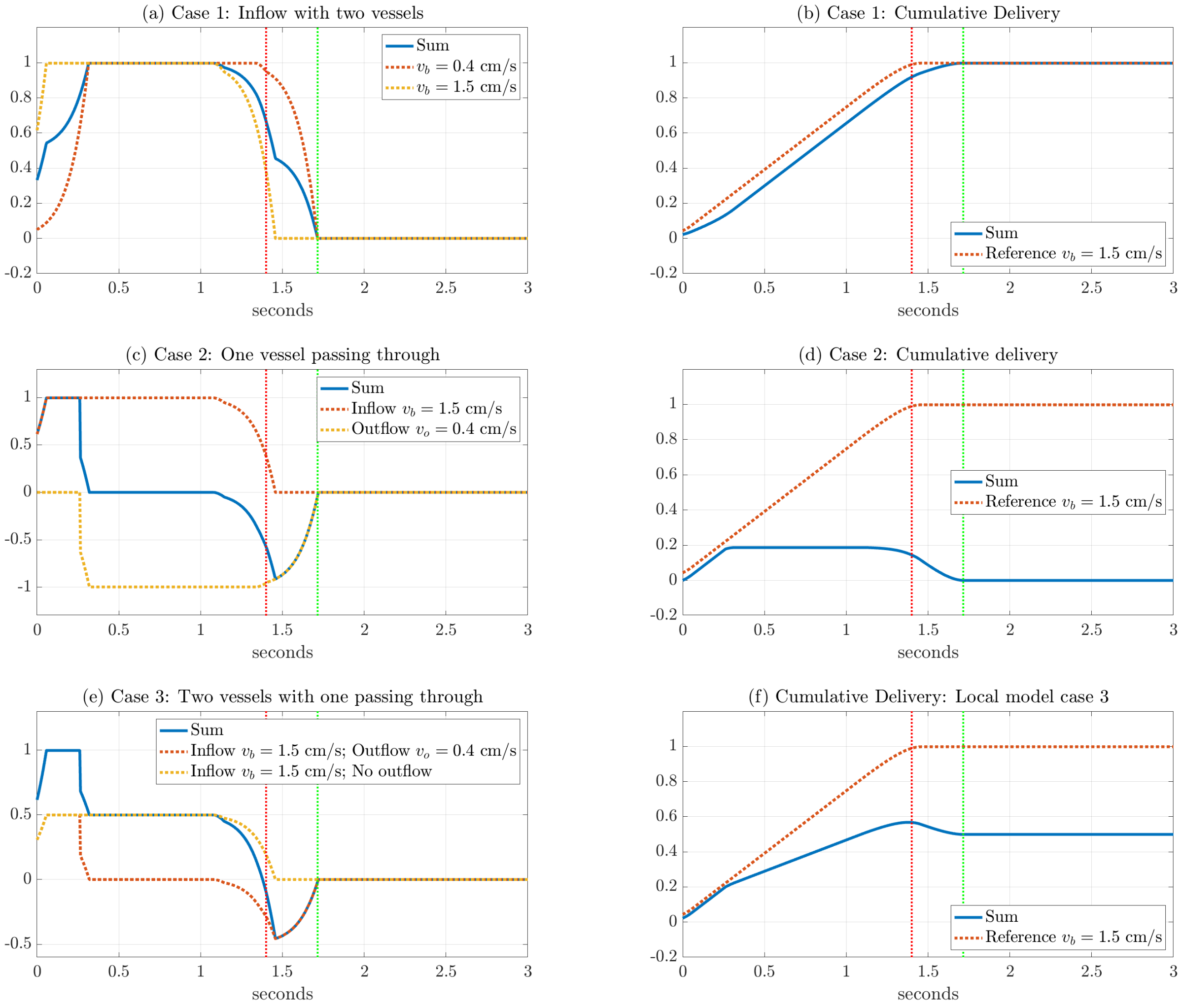


Figure S7: Arterial delivery functions (panels a, c, e) and cumulative delivery signals (panels b, d, f) for the local model case examples, assuming windowed cosine VSS with $v_c = 2$ cm/s. Note that the arterial delivery functions are normalized by $Q_0(\tilde{\mathbf{r}})$, and the cumulative delivery signals include the effect of an initial arterial volume term for vessels that deliver blood to the capillary bed. Case 1 (panels a, b): two arterioles with different boundary velocities (0.4 cm/s and 1.5 cm/s) deliver blood to the capillary bed. Case 2 (panels c, d): an arteriole passes through the voxel with entry and exit velocities of 1.5 cm/s and 0.4 cm/s, respectively. Case 3 (panels e, f): arteriole that branches into two child arterioles, with one delivering blood to the capillary bed and the other exiting the voxel. See text for additional details.

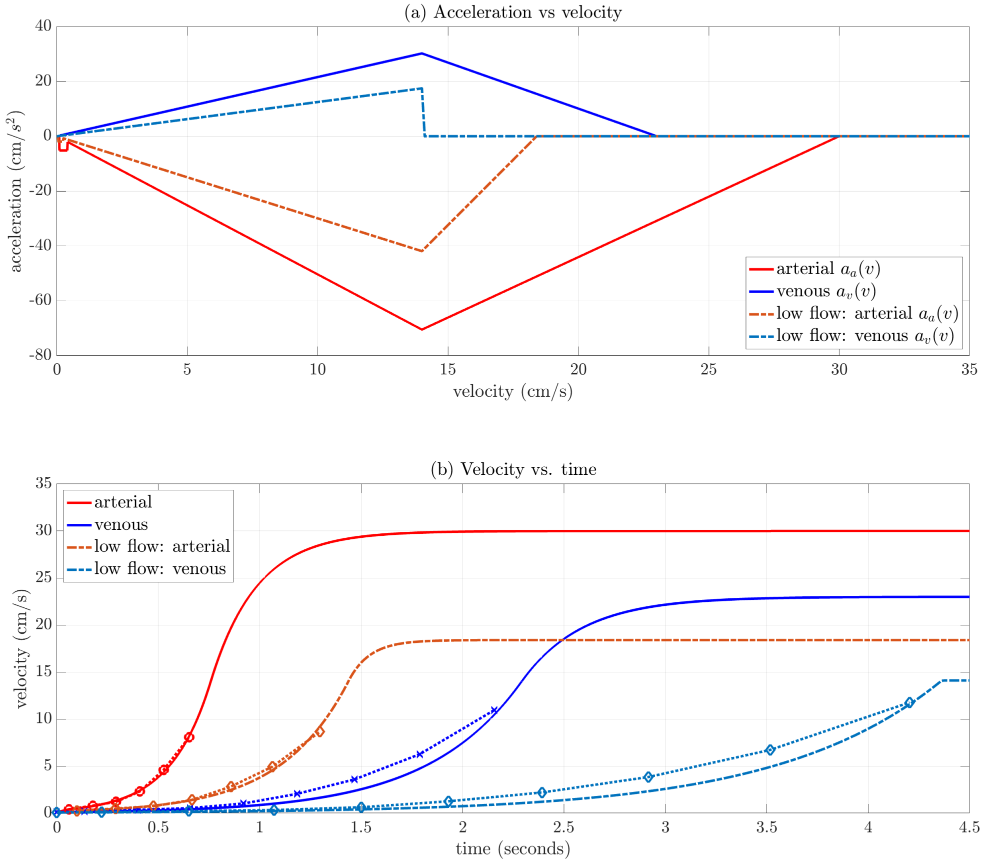


Figure S8: (a) Arterial $a_a(v)$ (red, brown) and venous $a_v(v)$ (blue, light blue) acceleration functions with parameters described in section S.1.1 for normal (solid) and low flow (dash) conditions. (b) Arterial $v_a(t, v_{cap})$ (red, brown) and venous $v_v(t, v_{cap})$ (blue, light blue) velocity as a function of time for normal (solid) and low flow (dash) conditions). For both panels, the Murray's law regime applies for velocities below 14 cm/s. In panel (b), symbols and dashed lines show values in this regime from the vascular model adapted from [8].

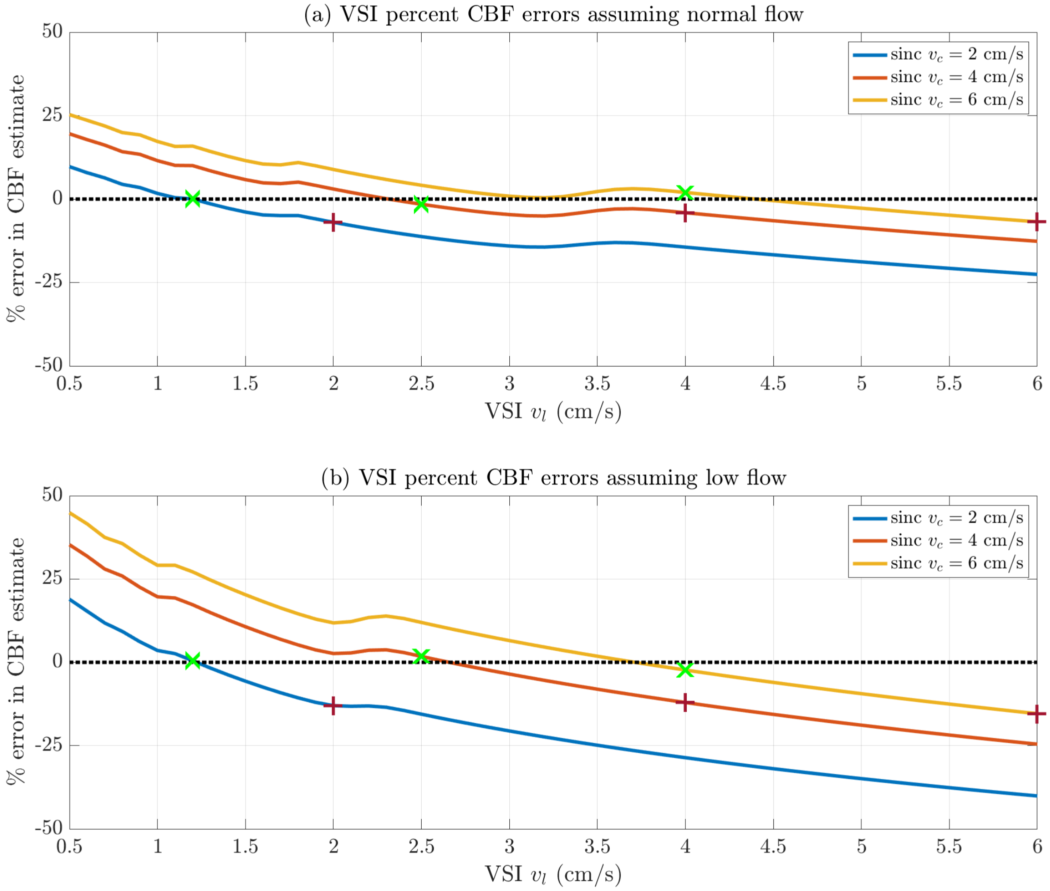


Figure S9: Percent error in CBF estimates for (a) normal flow and (b) low flow cases, obtained with Equation 24 using bolus width errors from Figure 8 and $\tau = 1.4$ s. Each curve shows the percent error as a function of FTVSI LCM cutoff velocity $v_l$ for a given value of sinc VCM cutoff velocity $v_c$. Errors obtained when $v_l = v_c$ are indicated by the red crosses. For each $v_c$ curve, the green x's indicate the error at optimal $v_l$ values that minimize the average error across normal and low flow conditions.

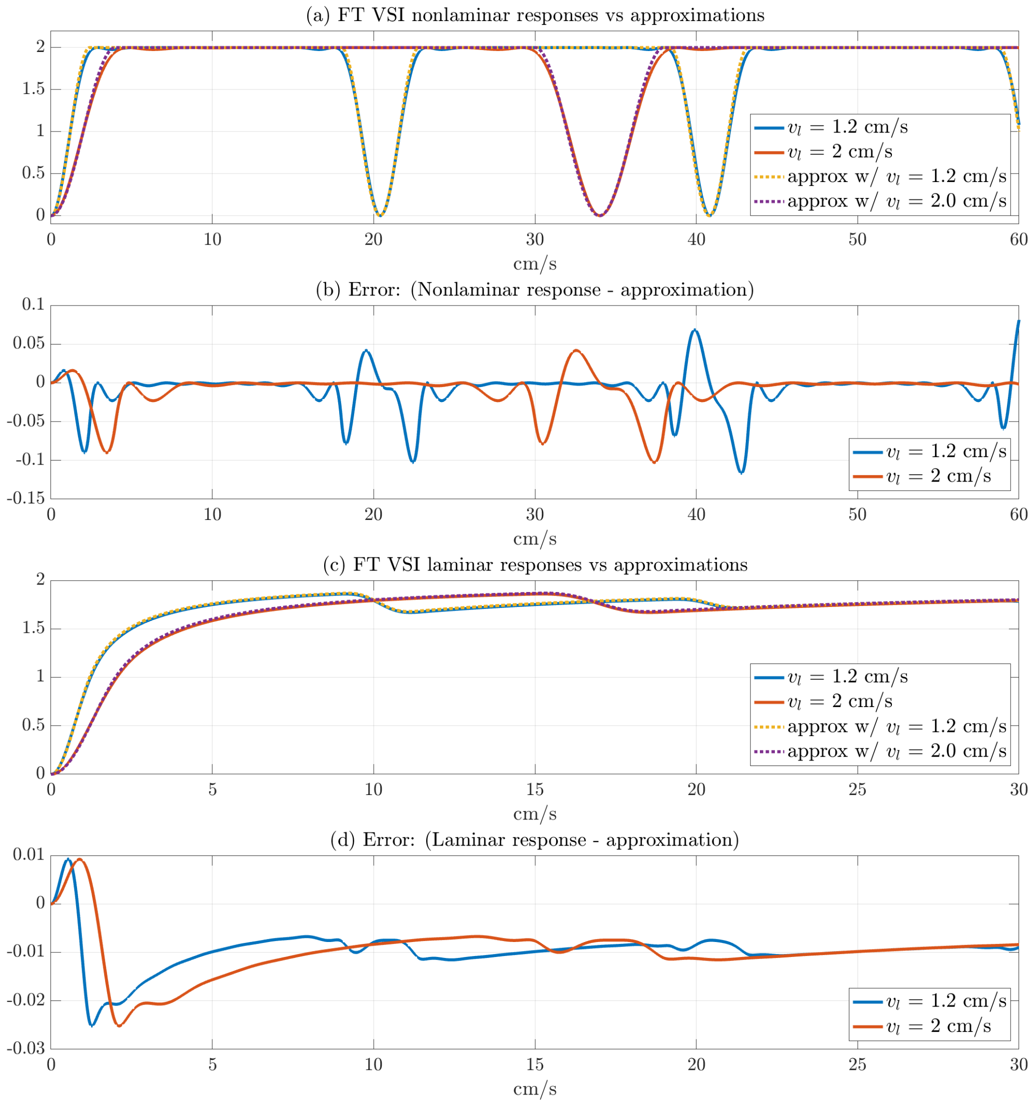


Figure S10: Approximations to the VSI passband functions. (a) Solid lines show Bloch simulated VSI passband functions for $v_l = 2.0$ cm/s and $v_l = 1.2$ cm/s prior to laminar flow integration, while the dotted lines show the corresponding approximations from Eq. A.19. (b) Differences between the simulated passband functions and the approximations. (c) Solid lines show Bloch simulated VSI passband functions after laminar flow integration, while dotted lines show corresponding approximations from Eq. A.20. (d) Differences between the simulated passband functions and the approximations with laminar flow integration. Note that the velocity range for the top two rows is double that of the bottom two rows, and the errors shown in (d) are an order of magnitude less than those in (b).
